# Molecular Diversity of Intrinsically Photosensitive Ganglion Cells

**DOI:** 10.1101/381004

**Authors:** Daniel Berg, Katherine Kartheiser, Megan Leyrer, Alexandra Saali, David Berson

**Affiliations:** Molecular Biology Program, Brown University, Providence, RI 02912; Department of Neuroscience, Brown University, Providence, RI 02912

## Abstract

Intrinsically photosensitive retinal ganglion cells (ipRGCs) are rare mammalian photoreceptors essential for non-image-forming vision functions, such as circadian photoentrainment and the pupillary light reflex. They comprise multiple subtypes distinguishable by morphology, physiology, projections, and levels of expression of melanopsin (Opn4), their photopigment. The molecular programs that differentiate ipRGCs from other ganglion cells and ipRGC subtypes from one another remain elusive. Here, we present comprehensive gene expression profiles of early postnatal and adult mouse ipRGCs purified from two lines of reporter mice marking different sets of ipRGC subtypes. We find dozens of novel genes highly enriched in ipRGCs. We reveal that Rasgrp1 and Tbx20 are selectively expressed in subsets of ipRGCs, though these molecularly defined groups imperfectly match established ipRGC subtypes. We demonstrate that the ipRGCs regulating circadian photoentrainment are unexpectedly diverse at the molecular level. Our findings reveal unexpected complexity in gene expression patterns across mammalian ipRGC subtypes.

## Introduction

Many unique attributes distinguish intrinsically photosensitive RGCs from conventional RGCs. Only ipRGCs express the blue-light sensitive photopigment melanopsin (OPN4), which renders them autonomously light-sensitive. They violate the usual stratification rule in which ON-type RGCs deploy their dendrites only in the inner (proximal) half of the inner plexiform layer; their inputs from ON bipolar cells are atypical (Dumitrescu et al., 2009; Hoshi et al., 2009; Kim et al., 2010). Whereas most RGCs direct their entire output through the optic nerve, some ipRGCs modulate intraretinal processing, through dopaminergic (Zhang et al., 2008; Xue et al., 2011) and other amacrine cells (Reifler et al., 2015; Sabbah et al., 2017) and spontaneous retinal waves during the early postnatal period (Renna et al., 2011). Additionally, ipRGCs appear more resistant than RGCs overall to various sorts of insults, including intraocular hypertension, optic nerve injury, glutamate-induced excitotoxicity, and glaucoma (Cui et al., 2015). Functionally, ipRGCs are unique among RGCs in their ability to encode overall light intensity for extended periods (Wong, 2012). This tonic luminance signal is transmitted to specific brain targets for a variety of functions. Projections to the hypothalamus mediate photoentrainment of circadian rhythms, while those to the midbrain to drive pupillary constriction. These outputs derive from molecularly distinct ipRGC subtypes.

The distinctive structural and functional properties of ipRGCs must ultimately be traceable to different patterns of gene expression. However, there is very little information on what these differences might be. For example, the basic molecular framework of the melanopsin phototransduction cascade, a major defining feature of ipRGCs, has only begun to be identified (Hughes et al., 2012) and the precise phototransduction mechanisms remain poorly characterized. More generally, very little is known about the developmental molecular mechanisms that direct certain immature RGCs to an ipRGC fate, or that maintain the distinctive features of ipRGCs throughout maturity. Thus, there is ample motivation for comparing the transcriptional profiles of ipRGCs as compared to those of conventional RGCs.

The ipRGCs consist of at least six anatomically distinct retinal subtypes, termed M1-M6. These which differ in their level of melanopsin expression, visual response properties, dendritic stratification, axonal projections, and contributions to light-modulated behavioral responses (Schmidt et al., 2011). Little is known about the gene regulatory programs that differentiate and maintain the specialized properties of individual ipRGC subtypes. M1 ipRGCs have been further subdivided based on their expression of the transcription factor Pou4f2 (Brn3b) (Chen et al., 2011; Jain et al., 2012). M1 ipRGCs that innervate the suprachiasmatic nucleus (SCN) do not express Brn3b, while those that project to other M1-cell targets, such as the olivary pretectal nucleus (OPN), do express this transcription factor (Chen et al., 2011). Indeed, ablation of Brn3b-positive ipRGCs severely impairs the pupillary light reflex, but leaves circadian photoentrainment intact (Chen et al., 2011). There is surely additional molecular diversity among ipRGCs, both within and between established ipRGC subtypes, and this further motivates the present study.

Previous attempts to develop a “molecular parts list” for ipRGCs through gene expression profiling of adult ipRGCs have been limited by the extreme heterogeneity of retinal tissue, and the fragility of mature retinal neurons, and the minuscule amount of genetic material from ipRGCs, which comprise far fewer than 1% of all retinal neurons (Lobo et al., 2006; Heiman et al., 2008; Sanes and Masland, 2015). Some progress has been made by purifying ipRGCs from enzymatically dissociated retinas using either anti-melanopsin immuno-panning or fluorescence-activated cell sorting (FACS) of genetically-labeled fluorescent ipRGCs. However, prior efforts have been limited by low yield and inclusion of contaminating cell populations such as rods (Hartwick et al., 2007; Peirson et al., 2007; Siegert et al., 2012).

Here we conducted a thorough unbiased transcriptomic analysis of ipRGCs by purifying GFP-tagged ipRGCs through a combination of FACS and immunoaffinity and comparing this the transcriptional profile of GFP-negative RGCs. We did this in two different mouse lines, marking partially overlapping subsets of ipRGCs. One was a BAC transgenic reporter (*Opn4-GFP*), which fluorescently labels M1-M3 ipRGCs (see Methods) and the other was a *Opn4-Cre;Z/EG* reporter system, which labels all ipRGC subtypes, M1-M6 (Schmidt et al., 2008; Ecker et al., 2010; Quattrochi et al., 2018; Stabio et al., 2018). The specificity and purity achieved by our approach is validated by the substantial enrichment in the ipRGC samples of known molecular markers of ipRGCs and by the fact that transcripts selectively expressed in potential contaminating cell types are generally absent. We identified more than 75 new gene candidates expressed much more highly in adult ipRGCs than in other RGCs. We validate two of the new molecular markers at the protein level: the Ras GEF Rasgrp1 and the T-box transcription factor Tbx20 and relate these to established ipRGC subtypes and patterns of central projection.

## Results

We enzymatically dissociated retinas from melanopsin-reporter mice, selected for RGCs by anti-Thy1 immunoaffinity, and sorted these into presumptive ipRGC and conventional ganglion-cell (cRGC) pools by FACS based on the fluorescent labeling of ipRGCs (see Methods, Figure 1A). We then compared gene expression in these ipRGC-enriched and cRGC-enriched cell samples to identify genes differentially expressed in ipRGCs as compared to other ganglion cells. We used two strains of melanopsin-reporter mice (see Methods). One of these was a BAC transgenic mouse (here termed *Opn4-GFP*) in which retinal GFP expression is apparently restricted to ipRGCs of subtypes M1, M2 and M3 (see Methods; personal communication R. Maloney and L. Quattrochi). This is presumably because expression of the reporter is coupled to expression of melanopsin, which is highest in this subset of ipRGCs. The other melanopsin reporter mouse was obtained by crossing a knock-in mouse in which *Cre* replaces *Opn4* (Opn4^cre/cre^) with a Cre reporter strain (Z/EG). The resulting Cre-driven labeling of melanopsin-expressing cells is more sensitive than in the other strain of reporter mice, and labels all known types of ipRGCs, M1-M6, while labeling few if any conventional RGCs (cRGCs) (Ecker et al., 2010; Estevez et al., 2012; Quattrochi et al., 2018; Stabio et al., 2018) (Figure 1A; see Methods).

**Figure 1.**
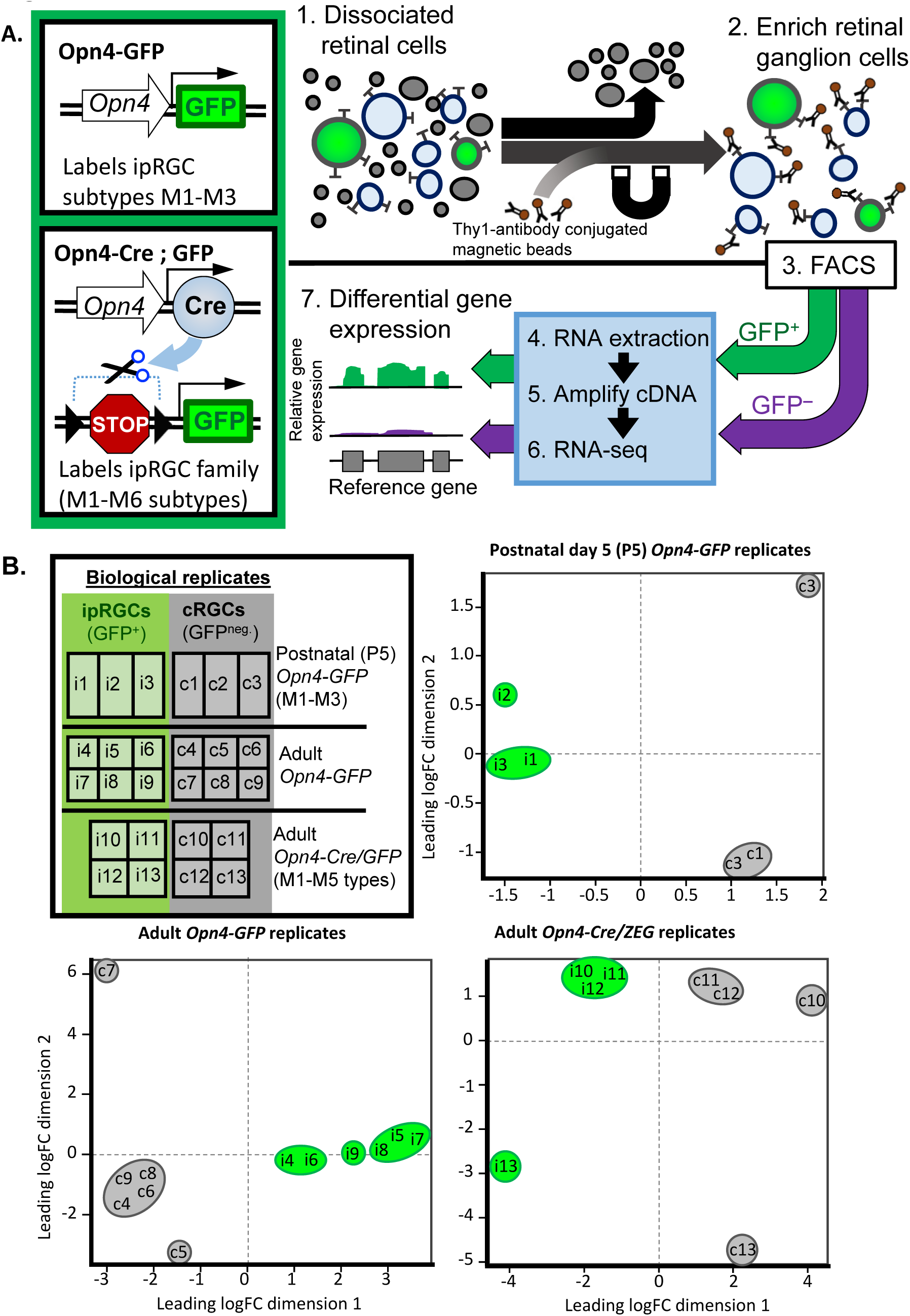
Experimental design of gene expression profiling from purified ipRGCs and comparison with generic RGCs. A. Two transgenic reporters were used for gene expression profiling of ipRGCs. The BAC transgenic Opn4-GFP labels M1-M3 ipRGCs while the Opn4-Cre crossed with a cre-dependent GFP reporter labels M1-M6 ipRGCs. B. Schematic of the gene expression profiling procedure. 1) Isolation of cell populations from enzymatically dissociated retinas. 2) The surface protein Thy-1 is enriched in RGCs, this high affinity of Thy1-conjugated magnetic beads to RGCs was used to enrich the extracted cell populations with RGCs. 3) Fluorescence-activated cell sorting (FACS) was used to isolate GFP-positive cells (ipRGCs) from GFP-negative cells (cRGCs). These two populations were isolated in parallel to provide direct internal testing of ipRGCs versus cRGCs under the same treatments, conditions, and genetic backgrounds. 4) The RNA of these two main populations was subjected to mRNA extraction. 5) The RNA was converted to cDNA and amplified using Nugen Ovation RNA amplification system. 6) Illumina Truseq sequencing libraries were prepared by ligating adapters to the cDNA. Single-end 50 base pair sequencing was completed using the Illumina HiSeq system. 7) Differentially expressed genes were determined using EdgeR bioinformatics pipeline. See Methods for details. B. EdgeR multi-dimensional scaling (MDS) plot illustrates the overall similarity between expression profiles of different samples. Each sample is denoted by a letter (“i” for ipRGCs; c for cRGCs) and a number, corresponding to particular replicate, comprising one pool of purified RGCs then divided into the two pools. Numbering scheme represents paired ipRGCs (GFP+) and cRGCs (GFP-) replicates (i.e., ipRGC sample ‘i1’ was processed in parallel with cRGC sample ‘c1’, sample ‘i2’ with ‘c2’, etc.). Distances are approximately the log2 fold changes between samples. Green and gray ovals represent ipRGC (GFP+) and cRGC (GFP-) samples, respectively. Adapted from EdgeR simple graphical output of individual samples in 2D space.

### Cell composition and purity of isolated ipRGCs and conventional RGCs

The relationship among the transcriptional profiles of ipRGC and cRGC samples across replicates are illustrated in the multidimensional scaling (MDS) plots of Fig. 1B. These show the relationship between all pairs of samples (one of ipRGCs, the other of cRGCs) based on a count-specific pairwise distance measure (Anders et al., 2013) (Figure 1B). These sample pairs were clearly separated along the first dimension, indicating pronounced differences in overall gene-expression patterns between ipRGC and cRGC samples. Samples of ipRGCs and cRGCs derived from the same retina and processed in parallel tended to be closely spaced along the second dimension, indicating greater similarity within than across replicates. This may reflect slight differences in overall genetic makeup of mice contributing to each pool, since both strains used were on a mixed genetic background, or to slight technical differences in the acquisition and processing of RNA from one run to the next.

Transcriptional data offer broad internal evidence for the efficacy of purification of cell samples. As expected, the *Opn4* (melanopsin) gene was among the genes much more highly expressed in ipRGCs than in cRGCs. For example, *Opn4* was enriched 40-fold in adult ipRGCs purified from *Opn4-GFP* mice, and this was highly significant, at q<1×10^−55^ false discovery rate (FDR). Though *Opn4* expression was detected in cRGCs at modest levels (Figure S1), this was expected, because some ipRGCs lack GFP expression in both melanopsin reporter lines (Opn4-GFP and Opn4-cre/GFP), and these would have been pooled with cRGCs during the FACS procedure.

Transcripts of other genes known to be expressed in ipRGCs were also enriched in the ipRGC pool relative to the cRGC pool (FDR < 0.05, significantly expressed in ipRGC samples, and absent or weakly expressed in cRGC samples). Among these genes were *Adcyap1* (pituitary adenylate-cyclase activating polypeptide; PACAP), *Tbr2*, (*Eomesodermin*), *Trpc7* and, to a lesser extent, *Trpc6* (Hannibal et al., 2004; Xue et al., 2011; Sand et al., 2012; Mao et al., 2014; Sweeney et al., 2014). Taken together, these results demonstrate that mRNA isolated from purified ipRGC samples were enriched as expected for transcripts for genes that are known to be differentially expressed in ipRGCs.

In the purified ipRGC samples, we found little or no evidence of contamination by transcripts from other retinal cell types. For example, transcript levels were very low for rod and cone opsins, for the amacrine-specific marker ChAT, for several bipolar markers (Otx2; Vsx2; Grm6; Trpm1), and for markers of astrocytes, microglial and vascular cells (Figure S1). Several transcripts suitable for assessing potential contamination from Müller glia (*Glul, Vim)* were present at surprisingly high levels in the purified ipRGC samples, suggesting that these glial cells may contaminate the transcriptional picture to some degree.

In general, the cRGC samples were relatively less pure than the ipRGCs samples by this measure. A particularly informative transcript for assessing such contamination is that for the rhodopsin gene (*Rho*), because rods are by far the most common neuronal type in the mouse retina and express *Rho at* very high levels. Rhodopsin transcripts were significantly (150-fold) more abundant in the cRGC samples than in ipRGC samples (Figure Supplement 1), whether isolated from *Opn4-GFP* or *Opn4-Cre/GFP* adult reporter mice (FDR < 6 x 10^−9^). Evidently, the second isolation step in which GFP+ positive cells were isolated by FACS from the purified RGC pool was a key factor in the greater purity of the ipRGC sample. Similarly, transcripts associated with bipolar cells and Müller glial cells were generally more abundant in cRGC than ipRGC samples. For example, the cRGCs had relatively high expression of the known Müller glia markers *Glul, Apoe, Aqp4*, and *Vim*, generally higher than in the ipRGC pool (Figure S1). Contamination of adult cRGC samples by other cell types may explain why most RGC markers, such as *Rbpms* and *Sncg* (Soto et al., 2008; Rodriguez et al., 2014), were less abundant in the cRGC cell pool than in the ipRGC pool. However, the data suggest that contamination in the cRGC pool was not uniform across retinal cell types. Amacrine-specific transcripts were no more abundant overall in cRGCs than in ipRGCs, and microglial and vascular markers were essentially absent, as in ipRGCs.

In immature mice (P5; *Opn4-GFP*), contamination of cRGC samples by non-RGC transcripts appeared more modest than in adults. The major sources of contamination (rods and Müller glia) are still being born and undergoing early-stage differentiation at this age, and this would presumably depress the abundance of their cell-type-specific transcripts (Young, 1985; Morrow et al., 1998; Matsushima et al., 2011).

To summarize, this analysis suggests that all samples were relatively free of contamination by most other retinal cell types, and that the ipRGC samples were particular pure. Contamination of the cRGC samples appears to derive mainly from Müller cells and strongly expressed genes in rods. Though this must be factored into the analysis, our primary focus was on genes more strongly expressed in ipRGCs than in cRGCs, and this difference seems unlikely to be affected by the modest contamination of the cRGC pool.

### Genes differentially expressed in ipRGCs

Comparing the abundance of transcripts in the ipRGC and cRGC pools, we identified over 75 genes that were differentially elevated expression in ipRGCs (as marked by one or both melanopsin-reporter lines) relative to cRGCs. Briefly, identification of differentially expressed genes in ipRGCs relied on the following stringent criteria: 1) low false discovery rate with high fold-change, 2) corroboration of differential expression across both ipRGC reporters, and 3) absence of gene expression in cRGC samples (see Methods). The identified differentially expressed genes are diverse, and most have not been previously identified as ipRGC-enriched (Figure 2; see Methods). Here, we survey some of these genes, grouped by their functional features (Figure 2).

**Figure 2.**
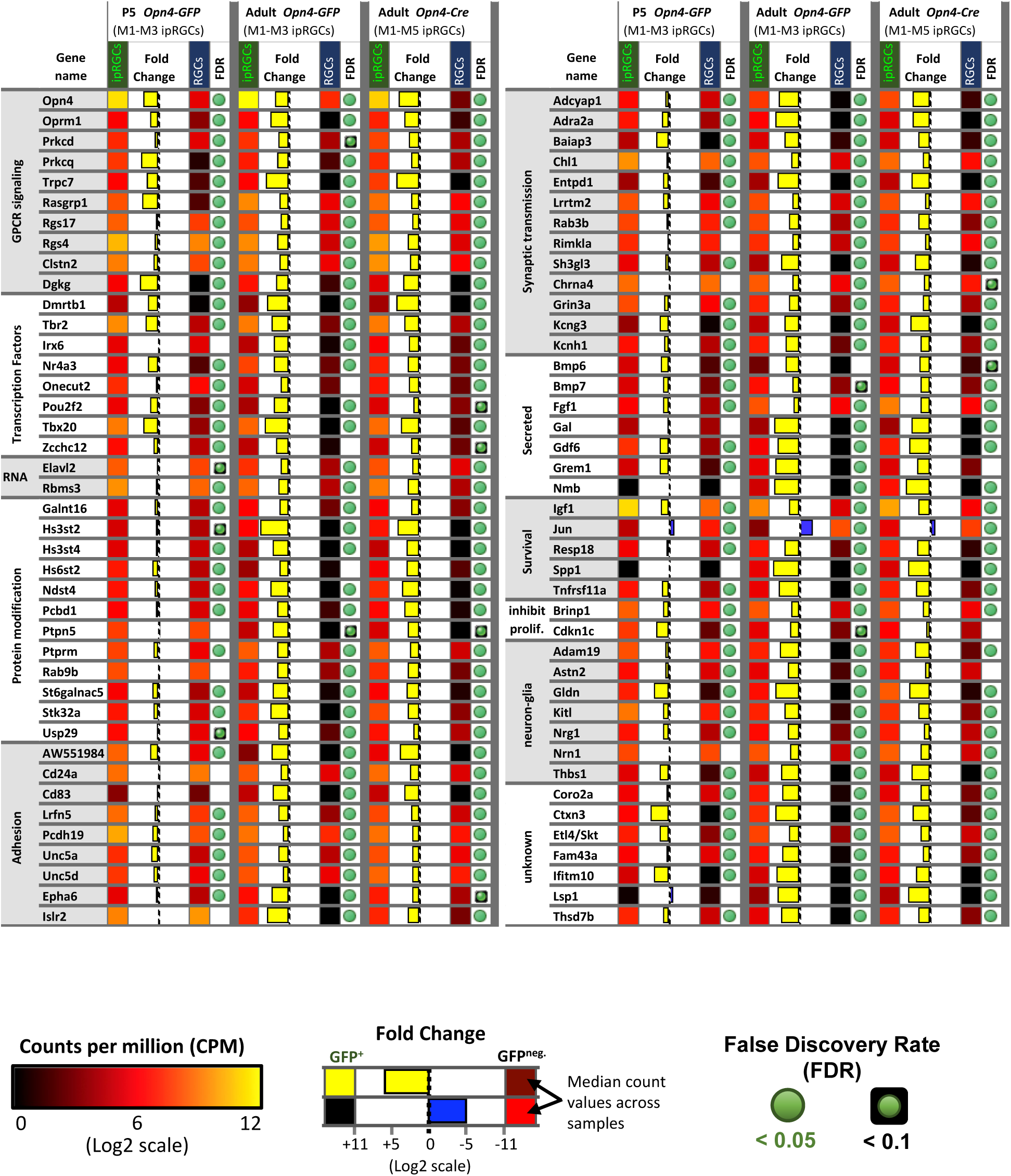
The expression pattern of candidate ipRGC-specific genes. Heat map of 83 genes differentially expressed in ipRGCs that have functional links to GPCR signaling, regulation and maintenance of molecular programs, neuron communication and organization, neuron survival, and neuron-glia interactions. Relative expression levels, fold change, and FDR are color-coded as indicated in the figure.

#### 1. Transcription factors

Transcription factors, by regulating other genes, help to generate and maintain ipRGC identity. We noted above that the T-box transcription factor *Tbr2* was much more strongly expressed in adult ipRGCs than in cRGCs, as expected (Sweeney et al., 2014). *Tbr2* is best known for its key role in early retinal development. Its expression in adult retina is far more restricted, but it remains expressed in the majority of ipRGCs. A second T-box transcription factor, *Tbx20*, was similarly enriched (Figure 2). *Tbx20* has not been previously linked to adult retinal function, but we will show that it too is quite selectively expressed in ipRGCs. Additionally, four other transcription factors, *Irx6, Dmrtb1, Nr4a3*, and *Pou6f2,* were differentially expressed in adult ipRGCs. Most of these genes serve as broad lineage determinants in early retinal development (Zhou et al., 1996; Star et al., 2012). Other highly expressed genes included the neuron-derived orphan receptor 1 *Nor1* (*Nr4a3*), which codes for a nuclear receptor, and *Elavl2* gene, which codes for a RNA-binding protein important for mRNA metabolism and neuronal differentiation (Fornaro et al., 2007; Hinman and Lou, 2008). Pathway analysis (DAVID) of differentially expressed genes in ipRGCs suggested specialization in heparan sulfate biosynthesis, including *Hs3st4, Hs3st2, Hs6st2, Ndst4*, and *Gpc5* (Figure 2).

#### 2. Synaptic transmission

Some of the genes differentially expressed in ipRGCs have known roles in regulating synaptic function, especially at presynaptic sites. Among others, these genes include *Sh3gl3, Entpd1, Rab3b, Baiap3, Chl1, Sh3gl3*, and *Adra2a* (Figure 2).

#### 3. Growth factors and neuropeptides

Multiple growth factors and neuropeptides were also differentially expressed in adult ipRGCs. These include *Bmp7, Fgf1, Gal, Gdf6, Grem1, Nmb* and *Nppb*) (Figure 2).

#### 4. Receptors and channels

Multiple genes encoding diverse surface receptors were differentially expressed in ipRGCs (Figure 2—figure supplement 2). For example, expression data suggest that ionotropic nicotinic acetylcholine receptors in ipRGCs may be composed of α3, α4, α6, β2 and β3 subunits (Figure 2—figure supplement 2), although the α3 and α4 transcripts were borderline for differential expression in ipRGCs. In agreement with previous studies, we found that ipRGCs expressed the Drd1 dopamine receptor, but had low levels of Drd2 expression (Van Hook et al., 2012). Several serotonin receptor genes (Htr1b, Htr1d, and Htr5a) were modestly enriched in ipRGCs. The ipRGCs were also found to express many glutamate receptors subunits, but only one of these - the NMDA receptor subunit 3A (GRIN3A) - was differentially expressed relative to other adult RGCs. The mu opioid receptor gene *Oprm1* is differentially expressed in ipRGCs; it could regulate their light responses interacting with L-type calcium channels, which carry the majority of light-evoked inward calcium current in ipRGCs (Moises et al., 1994; Diaz et al., 1995; Doğrul et al., 2001; Hartwick et al., 2007). Our data appear at odds with earlier reports that M1 and M4 ipRGCs express the melatonin receptor genes Mtnr1a and Mtnr1b (Sengupta et al., 2011; Pack et al., 2015; Sheng et al., 2015). Additionally, *Kcnh1*, also known as ether-a-go-go (*Eag1*), was differentially expressed in ipRGCs (Figure 2). Kcnh1 is a voltage-gated K+ channel that has been shown to be crucial for the generation of dark current in the inner segment of rods (Frings et al., 1998), but may normally regulate other neuronal functions in ipRGCs (Martin et al., 2008).

#### 5. Cell adhesion

Genes encoding for several cell adhesion molecules were differentially expressed in ipRGCs (Figure 2—figure supplement 3B). For example, the cell adhesion molecule *DscamL1* was relatively low in ipRGCs during postnatal development, but the closely related genes encoding the immunoglobulin superfamily (IgSF) adhesion molecules *Sidekick-1* and *Sidekick-2* were enriched in developing ipRGCs. *Unc5a* and *Unc5d* were significantly differentially expressed both in early postnatal and adult ipRGCs. In contrast, expression of *Unc5b* and *Unc5C* in ipRGCs was low relative to that in cRGCs. As suggested previously, expression of the repulsive ligand *Sema6a* was significantly lower in ipRGCs than cRGCs during postnatal development (Matsuoka et al., 2011). However, its receptor *PlxnA4* was enriched in P5 ipRGCs. Another semaphorin, *Sema5a*, was also significantly enriched in developing ipRGCs. Other differentially expressed cell-adhesion molecules *Salm5* (*Lrfn5*), *Clstn2, Thbs1, Lrrtm2, Pcdh19, Ptprm*, and *Lrrc4c (Ngl1)* could play significant roles in the formation of ipRGC synapses (Burden-Gulley and Brady-Kalnay, 1999; Lin et al., 2003; de Wit et al., 2009; Xu et al., 2010; Lipina et al., 2016; Pederick et al., 2016; Lin et al., 2018). The cell surface glycoprotein *Mdga1* was also differentially expressed in developing ipRGCs, and is known to influence the formation and maintenance of inhibitory synapses (Pettem et al., 2013).

#### 6. Tolerance to stress

There is increasing evidence that ipRGCs are resistant to stress and able to survive under circumstances that are fatal for other retinal neurons (Li et al., 2008; de Sevilla Müller et al., 2014; Cui et al., 2015; Duan et al., 2015). The harsh dissociation and FACS processing has the potential of generating stress-induced gene expression changes (Figure 1 and 2, see methods). We attempted to identify potential survival molecular programs that are specific to ipRGCs compared to generic RGCs. The genes *Adcyap1* (PACAP), *Igf1*, and *Spp1* (*osteopontin*), all of which have previously described roles in promoting ipRGC survival (Atlasz et al., 2010; Duan et al., 2015) were differentially expressed in ipRGCs. We also identified a number of genes related to glial function differentially expressed in ipRGCs, including *Gldn, Cntn2, Lama4*, and *Astn2*, and *Thbs1* (Figure 2).

#### 7. Phototransduction

Photoactivation of melanopsin photopigment typically triggers a phosphoinositide signaling cascade resembling that in rhabdomeric (invertebrate) photoreceptors, involving G proteins in the Gq family, phospholipase C, and canonical TRP channels. In ipRGCs, the phototransduction cascade typically signals through Gq-family proteins and phospholipase C beta 4 (PLCB4) to open canonical TRP channels (Trpc7 and Trpc6) (Graham et al., 2008; Xue et al., 2011; Hu et al., 2013; Emanuel and Do, 2015; Emanuel et al., 2017) (Figure 2—figure supplement 4A). We determined that the genes in this signaling cascade (*Opn4, Trpc7, Trpc6, Plcb4*, and several Gq genes) were expressed at relatively high levels in all three ipRGC pools (i.e., selective-postnatal; selective-adult; or pan-subtype adult). Moreover, two key genes - *Opn4* and *Trpc7* - were more highly expressed in ipRGCs than in cRGCs in all three ipRGC pools. *Trpc6* was also significantly overexpressed in ipRGCs in younger animals, with a trend in this direction also in adult ipRGCs, but *Trpc7* was expressed at much higher levels than *Trpc6*. Plcb4 appears essential for melanopsin phototransduction in some cells, and it was expressed at much higher levels than Plcb1, 2 or 3. However, except in young mice, it was not more highly expressed in ipRGCs than cRGCs.

Only recently has the precise identity of the Gα subunits combination in ipRGCs been identified as redundantly expressing and signaling through the Gnaq, Gna11, or Gna14 subunits (Hughes et al., 2015). Our studies suggest a similar expression pattern, including a lack of Gna15 expression (Figure 2—figure supplement 4A). Further, we determined that Gna14 was differentially expressed in our P5 ipRGC samples, but it did not reach a statistical significant difference in adult Opn4-GFP ipRGCs. Gnaq appears to be among the highest expressing Gq/11 subunits in our study, in contrast to the lack of Gnaq gene expression detected by Siegert and colleagues (2012). To date, the Gβγ complex remains unknown. Our studies determined that the beta subunit Gnb1 is by far the highest expressing, having a 15-fold higher expression than the other subunits Gnb2, Gnb4, or Gnb5; while Gnb3 showed no expression in adult ipRGCs (Figure 2—figure supplement 4C). Additionally, we found that the gamma subunit Gng4 is differentially expressed in ipRGCs.

Also differentially expressed in ipRGCs were two factors with known roles in diacylglycerol (DAG) signaling, *Rasgrp1* and *Dgkg* (Figure 2—figure supplement 4B). Ras guanyl nucleotide-releasing protein 1 (*Rasgrp1*) is a guanine nucleotide exchange factor (GEF) that activates *Ras* by facilitating its GTP binding (Bivona et al., 2003). Rasgrp1 binds DAG and Ca^2+^, both of which are elevated by melanopsin phototransduction. This provides a possible basis for intrinsic photoresponses of ipRGCs to modulate Ras signaling and thus genes governing cell growth, differentiation and survival. We will return to a more detailed consideration of Rasgrp1 later in this report. Diacylglycerol kinase gamma (*Dgkg*) (Bivona et al., 2003; Shulga et al., 2011) converts DAG to phosphatidic acid, thus acting as a terminator of DAG signaling. Because DAG appears to be a key link between early steps in phototransduction and gating of the light-activated channels, Dgkg may regulate the kinetics of the photoresponse in ipRGCs. The protein products of the two overexpressed genes may interact. Diacylglycerol kinases are also known to bind to Rasgrp and modulate its activity (Topham and Prescott, 2001). Diacylglycerol and calcium are also known to activate the protein kinase C (PKC) family members Prkcd and Prkcq (Oancea and Meyer, 1998), which we determined to be differentially expressed in ipRGCs. Protein kinase C (PKC) activity has been suggested to be important for deactivating TRPC activity in the invertebrate photoreceptors and potentially also for the Opn4 phototransduction cascade (Graham et al., 2008). Peirson and colleagues (2007) previously identified another PKC member, Prkcz, as being important for ipRGC-mediated photoentrainment of circadian rhythms (Peirson et al., 2007). However, Prkcz is only moderately expressed and similar to control samples in our study.

In other photoreceptors, RGS (Regulator of G protein signaling) proteins play a key role in terminating the photoresponse by accelerating the intrinsic GTPase activity of the cognate G-protein (e.g., transducin in rods). Two RGS genes were overexpressed in all three ipRGC pools: *Rgs4* and *Rgs17*. At least one of these (Rgs17) regulates Gq signaling (Mao et al., 2004; Ji et al., 2011) (Figure 2—figure supplement 4B).

The arrestins also contribute to response termination by binding to phosphorylated opsin. ipRGCs exhibited strong expression of both beta arrestin genes (Arrb1, Arrb2) but low expression of rod (*Sag*) and cone (*Arr3*) arrestin genes. This is consistent with earlier evidence that beta arrestins rather than conventional retinal arrestins bind photoactivated melanopsin in ipRGCs (Cameron and Robinson, 2014). Still, these beta arrestin transcripts are both at similarly high levels in cRGCs as in ipRGCs, presumably because these arrestins regulate diverse GPCRs (Figure 2—figure supplement 4B).

Many of the genes involved in rod and cone phototransduction had low expression (scarce or no read alignment) and/or were present at much lower levels in ipRGCs than cRGCs. These include the genes for opsins, transducin alpha, and arrestin in rods (*Rho, Gnat1, Sag*) and cones (*Opn1mw, Opn1sw, Gnat2,* and *Arr3*; Figure 2—figure supplement 4C). Although Cngb1 was differentially expressed in ipRGCs, the total reads aligning to the Cngb1 locus were low and derived mainly from a limited region of the gene, and the obligatory alpha subunits were not detected, so this could be a false positive (Figure 2—figure supplement 4C).

### Genes differentially repressed in ipRGCs

The lack of contamination by non-RGC retinal neurons in the postnatal day 5 (P5) samples allowed us to identify genes that were differentially repressed in ipRGCs compared to cRGCs in early postnatal development. Our data suggested that the transcription factor *Jun* (Jun Proto-Oncogene) and *Irx4* are differentially repressed in P5 ipRGCs samples (Figure 2—figure supplement 3). Other genes that were differentially repressed in the P5 ipRGC samples included *Satb1, Satb2*, and *Foxp2* showed, all of which are known to have restricted expression in the abundant F-RGC type that is likely included in the cRGC samples (Rousso et al., 2016). The *Pou4f1* (*Brn3a)* and *Pou4f3* (*Brn3c*) transcription factors were both differentially repressed in P5 ipRGCs, consistent with their known lack of expression in ipRGCs (Jain et al., 2012) (Figure 2— figure supplement 3). The transcriptional repressors *Bcl11b (CTIP2), Irx4*, and *Tbr1* were all found to be differentially repressed in ipRGC compared to cRGCs samples. Furthermore, the *Cdkn1c* (*p57KIP2),* a gene known to be transcriptionally repressed by Bcl11b (Topark-Ngarm et al., 2006), had relatively increased expression in ipRGCs (Figure 2—figure supplement 3).

### Expression differences between the *Opn4-GFP* and *Opn4-Cre/GFP* reporter systems

To study gene expression differences across the different ipRGC subtypes, we compared the expression patterns of *Opn4-Cre/GFP* (labels all M1-M5 subtypes) and *Opn4-GFP* (labels only the M1-M3 subtypes) (Figure 2—figure supplement 1). In general, genes differentially expressed in ipRGCs identified in the two reporter systems were both supportive and cross-correlated. However, we identified 24 genes that were differentially expressed in the adult *Opn4-Cre/GFP* reporter but had low or no apparent expression in the *Opn4-GFP* reporter, suggesting selective expression in one or more of the M4-M6 ipRGC subtypes. The *Opn4-Cre/GFP* specific genes included *Anxa2, Gem, Sema3d, Rbp4*, and *Rxrg*. Recently, an Rpb4 reporter (Rbp4-Cre) was demonstrated to mark amacrine cells coupled to ipRGCs, although there was an apparent lack of labeling in ipRGCs (Sabbah et al., 2017). The Kcnk4/TRAAK, another gene that was differentially expressed in the *Opn4-Cre/GFP* reporter, encodes a two-pore potassium channel subunit (Fink et al., 1998). Additionally, our data suggest that the Kcns3 electrically silent voltage-gated potassium channel subunit has its expression restricted to the ipRGCs labeled by *Opn4-Cre/GFP*, but this difference did not reach statistical significance (FDR 0.13). However, close inspection of reads aligning to Kcns3 using the integrated genome viewer (IGV) confirmed weak expression in ipRGCs and absent expression in cRGCs for *Opn4-Cre/GFP* samples (data not shown). Lastly, the neurexophilins Nxph1 and Nxph3 were differentially expressed in the *Opn4-GFP* and *Opn4-Cre/GFP* reporters, respectively (Figure 2—figure supplement 1). These proteins are known to bind α-neurexins in mice and have restricted expression patterns (Missler et al., 1998; Beglopoulos et al., 2005; Craig and Kang, 2007).

### Rasgrp1 is selectively expressed in ipRGCs

We sought to test our transcript-level differential expression analysis at the protein level and to determine whether their expression is selective for particular adult ipRGC subtypes. Transcriptional profiling suggested that Rasgrp1 is expressed differentially, possibly even selectively, in ipRGCs. However, differential mRNA expression does not guarantee a correspondence with protein product (Koussounadis et al., 2015). Therefore, we used immunofluorescence against Rasgrp1 (Puente et al., 2000) to label the Rasgrp1 protein in whole-mount retinas from adult wild type mice. Rasgrp1-immunopositive somata were present in the ganglion cell layer (GCL) and in the inner nuclear layer (INL). The latter likely represent amacrine cells or displaced ganglion cells, judging by their close proximity to the inner plexiform layer (IPL) (Figure 3). Immunolabeling marked the cytoplasm as well as the somatic plasma membrane of these cells. Occasionally, particularly strongly Rasgrp1-labeled cells had some dendritic labeling. Rasgrp1 immunostaining was also observed in a subset of photoreceptors in the outer retina (data not shown).

**Figure 3.**
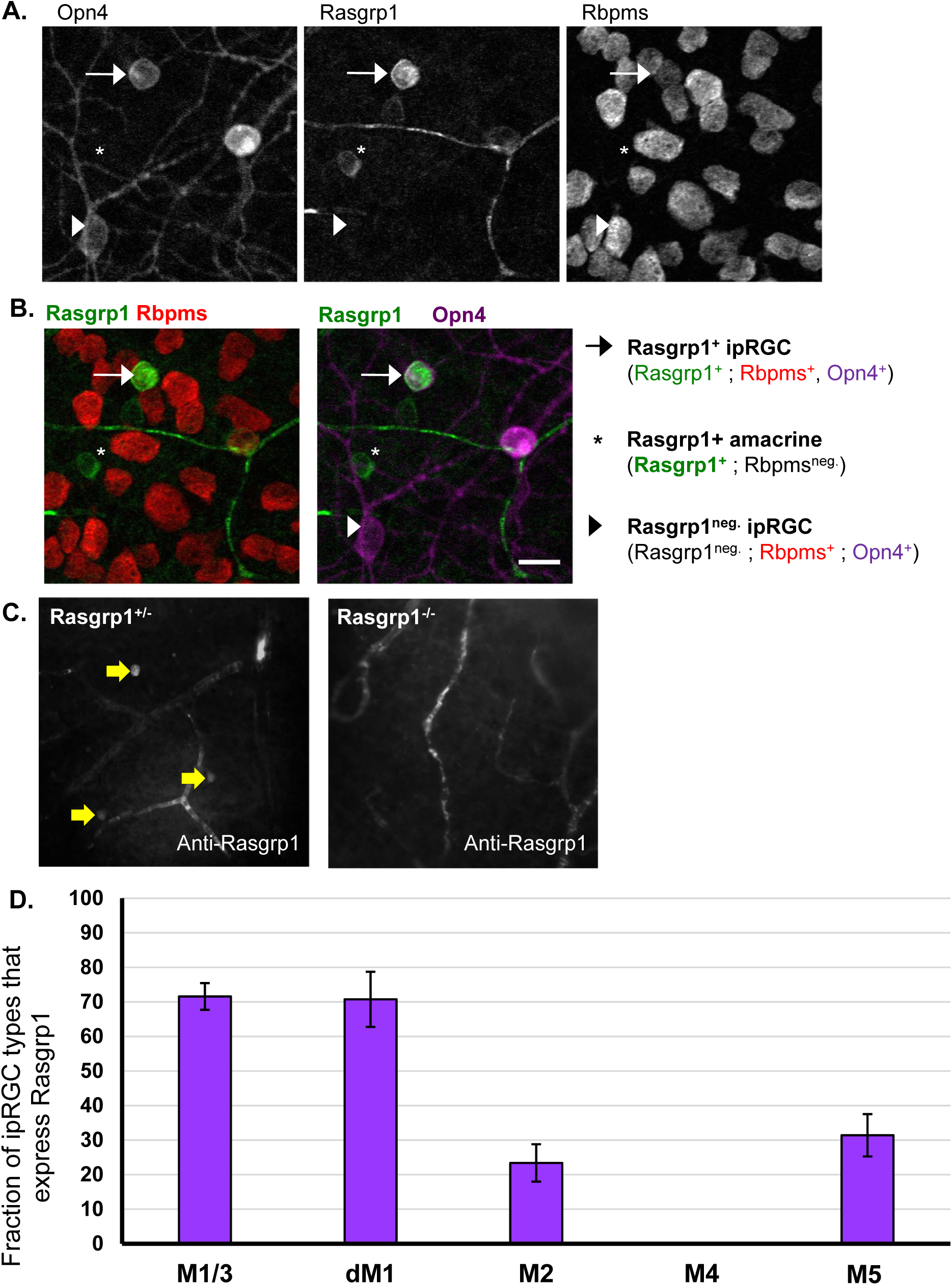
Rasgrp1 is selectively expressed in ipRGCs. A. Whole mount retina immunostained for Opn4, Rasgrp1, and the pan-RGC marker Rbpms (gray-scale). Focal plane is at the ganglion cell layer (GCL). We quantified co-localization of the three markers in confocal images of 49 regions that were topographically dispersed across three whole-mount adult retinas. B. Co-localization of Rasgrp1 (green), Rbpms (red), and Opn4 (magenta). Rasgrp1 is expressed in a subpopulation of amacrine cells and RGCs (Rbpms-negative and -positive, respectively). Scale bar, 20 µm. C. Rasgrp1 immunolabeling (antibody sc-8430) of cell bodies in GCL of Rasgrp1^+/-^ heterozygous mice (left, yellow arrows). Absence of cell body immunolabeling in Rasgrp1^−/−^ knockout mice (right) suggests a lack of cellular off-target antibody staining. D. Quantification of Rasgrp1-expression across Opn4-immunopositive ipRGC subtypes. 70% of M1 cells were Rasgrp1 immunopositive while only 20-30% of either M2, M5 or M6 cells were Rasgrp1 immunopositive. None of the identified M4 cells were Rasgrp1 immunopositive. M1 and M3 types were combined during the process of co-expression analysis (designated “M1/M3”). Error bars represent standard error of the mean.

To test whether the Rasgrp1-positive cells in the ganglion-cell layer were RGCs, we carried out double immunofluorescence for both Rasgrp1 (antibody m199) and the RNA-binding protein Rbpms, which selectively labels all and only RGCs (Rodriguez et al., 2014). About half of Rasgrp1-immunopositive cells in the GCL were RGCs, as determined by co-labeling for Rbpms (56%, *n* = 708 across three retinas, Figure 3A,B). The remainder can be assumed to be displaced amacrine cells.

Most of these Rasgrp1-expressing RGCs were ipRGCs, as revealed by their immunoreactivity for melanopsin (95.9±1.1%, *n* = 412, Figure 3A,B). In contrast, only a fraction of Opn4-immunopositive ipRGCs were Rasgrp1 immunopositive (34%, *n* = 1169). Thus, Rasgrp1 expression in the GCL is apparently restricted to a subpopulation of ipRGCs.

We next tested whether the immunolabeling of RGCs represented endogenous Rasgrp1 protein expression. The antibody used in this study has been previously shown to specifically label Rasgrp1 protein expression in hippocampal neurons (Pierret et al., 2000). As a further test for the specificity of the antibody, we compared immunofluorescence labelling of whole mount retinas from normal and Rasgrp1-knockout mice generated by inserting *LacZ* and a *Neo* cassette into the Rasgrp1 gene to disrupt its expression (Dower et al., 2000). Our control experiments showed that the GCL and INL cellular immunolabeling is absent in the Rasgrp1 knockout (Figure 3C). However, vasculature and photoreceptor cell labeling persisted in Rasgrp1-knockout mouse retinas, suggesting cross-reactivity of antibody with other proteins.

### Rasgrp1 expression is restricted to diverse ipRGC subtypes

We next determined which of the established morphological subtypes of ipRGCs express Rasgrp1 into adulthood (Figure 3D). For this purpose, we used key characteristics such as relative Opn4 expression, soma size, and dendritic morphology (see Methods). In the GCL, the majority of M1/3 cells (71.6±3.9%, *n* = 300 across three retinas) but only a fraction of M2 cells (23.4±5.4%, *n* = 389) and M5/6 cells (31.4±6.13%, *n* = 138) expressed Rasgrp1 (Figure 3D). In the INL, displaced M1 cells express Rasgrp1 at a similar percentage as conventionally placed M1 cells (70.7±8.0%, *n* = 3 retinas). We found no examples of Rasgrp1 immunoreactivity in M4 cells (0%, *n* = 172, Figure 3D). Of the Rasgrp1-expressing ipRGCs, half were M1/3 cells (52.8±3.8%), nearly a quarter were M2 cells (21.1±2.7%) and a small percent (10.4±1.8%) were M5/6 cells (*n* = 396 Rasgrp1^+^/Opn4^+^ cells across three retinas). Therefore, Rasgrp1 is selectively expressed in a diverse set of ipRGC subtypes.

### Spatial Distribution of Rasgrp1-expressing ipRGCs and amacrine cells

Our data show neither a ventral-dorsal nor a naso-temporal density gradient of Rasgrp1-expressing M1-M3 ipRGC across the retina (Figure 3—figure supplement 1C). However, we did observe a minor ventral-dorsal gradient of Rasgrp1-expressing RGCs, with a higher density in the ventral (65%) compared to the dorsal (35%) retina (Figure 3—figure supplement 1E). As shown above, Rasgrp1-expressing RGCs are almost exclusively ipRGCs (96%).

### Tbx20 is expressed in a diverse subset of ipRGCs

The T-box transcription factor Tbx20 was suggested from our gene expression analysis to be differentially expressed in ipRGCs. Immunofluorescence co-loclalization analysis of Tbx20 and Opn4 expression confirmed its high expression in a subset of ipRGCs (Figure 4D). Tbx20 was expressed in most M1 cells (82.6±1.8%, n=514 across four retinas), but only in a minority of M2/3 cells (30.2±6.5%, n=1305) and M5/6 cells (12.4±3.4%, *n* = 603). Half of the displaced M1 (dM1) cells expressed Tbx20 (46.0±7.1%, *n* = 153). Strikingly, however, Tbx20 was not expressed in M4 cells (0%, *n* = 283). These results demonstrate that Tbx20 is expressed in a diverse subset of ipRGCs.

**Figure 4.**
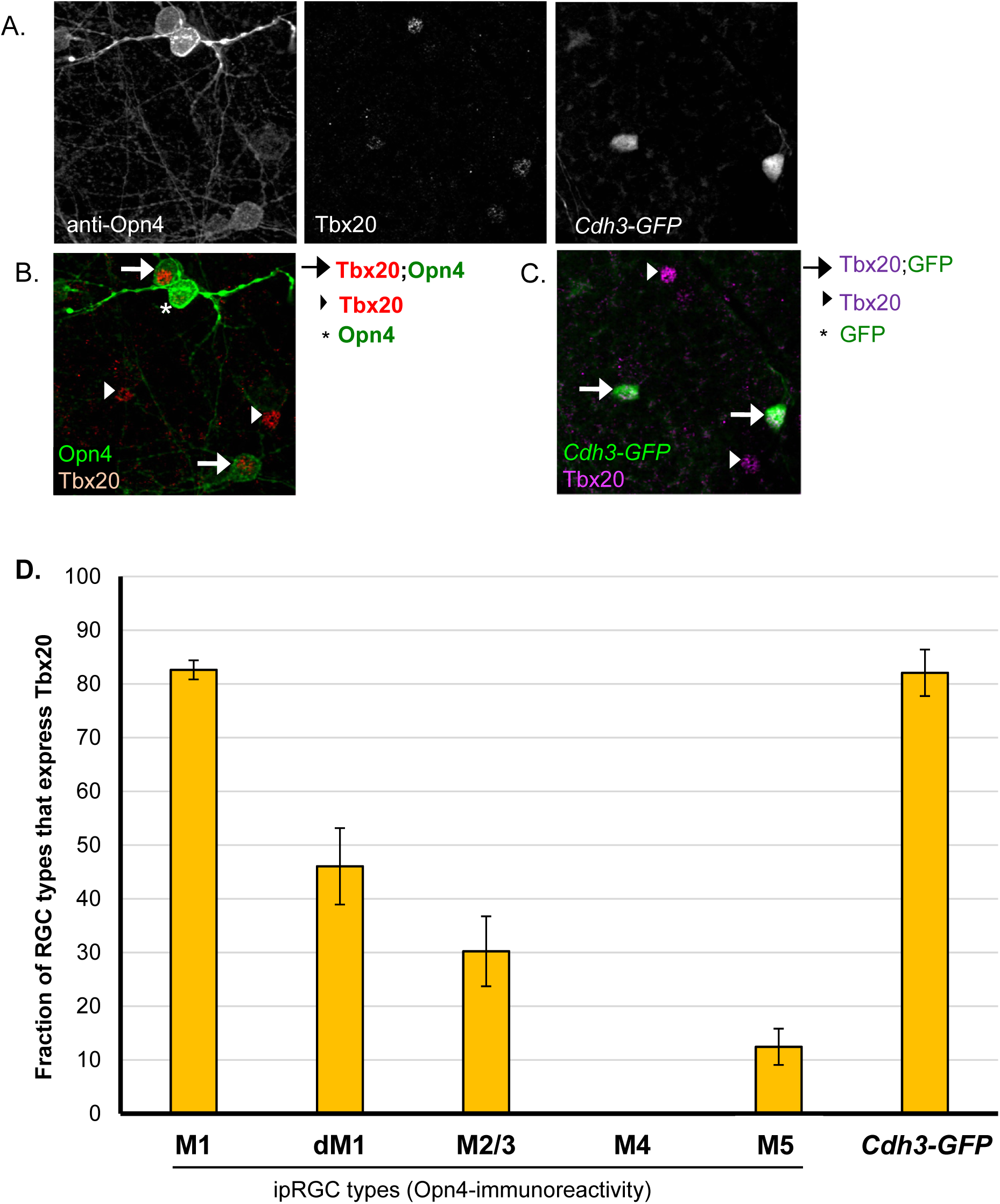
Colocalization study of Tbx20-expression in ipRGC subtypes. A. Triple immunofluorescence of Opn4, Tbx20, and *Cdh3-GFP* (gray scale). B. Tbx20-expression in subset of M1-M3 ipRGCs as well as an additional population of Opn4-immunonegative cells. B,C. Tbx20 is concentrated in the nucleus of most *Cdh3-GFP*-cells. D. Quantification of Tbx20-expression across Opn4-immunopositive ipRGC subtypes. Tbx20 immunofluorescence labels multiple ipRGC subtypes, including M1s, M2 cells and small soma, low Opn4 expression cells (presumptive M5/6 ipRGCs), and Cdh3-GFP cells (M6-type enriched). M2 and M3 types were combined during the process of co-expression analysis (designated “M2/M3”). Error bars represent standard error of the mean.

Many Tbx20 cells were not detectably immunopositive for Opn4. Only 41% of Tbx20 immunopositive were also Opn4-immunoreactive (18.5±2.6% were M1 cells; 16.0±2.0% were M2/3 cells; and only 3.6±0.9% were M5/6 cells; *n* = 2184 across four retinas; Figure 4—figure supplement 1). The remaining Tbx20-immunopositive cells were RGCs, as confirmed by Rbpms-immunoreactivity (data not shown). Additionally, Tbx20-immunopositive RGCs that were also Opn4-immunonegative were topographically enriched in the ventral retina, with most Tbx20-positive cells in the dorsal retina being accounted for by Opn4-immunoreactivity.

### Tbx20 expression in M5-M6 ipRGCs

To investigate whether some or all of the Tbx20-immunopositive RGCs that were Opn4-immunonegative might be ipRGCs of the M5 and M6 subtypes that exhibit weak Opn4 immunostaining, we examined the co-localization of Tbx20-immunopositive cells, Opn4-immunopositive cells, and all GFP-labeled cells in the *Opn4-Cre;Z/EG* mouse reporter, which among other ipRGCs, labels M5 and M6 cells. We observed examples of Tbx20-immunopositive cells that were GFP-positive (M1-M6 ipRGCs), but not Opn4-immunopositive (M1-M4), suggesting that Tbx20 is expressed in at least a subset of M5 or M6 ipRGCs (Figure 4—figure supplement 1A,B).

To test the implication that many Tbx20 cells were M5 or M6, we turned to Cdh3-GFP mice. Recently, our group determined that essentially all GFP+ RGCs in the Cdh3-GFP reporter mouse you used in your studies are either M6 cells (the great majority) or M5 cells (minority) (Quattrochi et al., 2018). We tested Tbx20 immunoreactivity in the context of Opn4 immunofluorescence and *Cdh3-GFP* labeling (Figure 4B,C). For the purpose of this study, we focused on GFP cells in the GCL that are Opn4-immunonegative (to distinguish from Opn4-immunopositive M2 types). We found that at three weeks after birth, most *Cdh3-GFP* cells express the Tbx20 protein (82.1±4.3%, *n* = 439 across four retinas, Figure 4C). Many, but not all, of the Opn4-immunonegative Tbx20-positive cells were GFP^+^ (27%, n=1277 Tbx20^+^;Opn4^−^ cells; 4 retinas). The dorsal-ventral gradient of Tbx20-positive cells that are Opn4-immunonegative was broadly similar to the retinal labeling of the *Cdh3-GFP* reporter. A large portion of Tbx20-immunopositive cells remained unclassified (43.3±3.8%, n=2184; Figure 4—figure supplement 1D).

Further, we determined whether Tbx20 expression correlates with the related T-box transcription factor Tbr2, a gene previously described to be enriched in adult ipRGCs (Mao et al., 2014; Sweeney et al., 2014). All Tbx20-expressing cells were strongly Tbr2-immunopositive (*n* = 328; 4 regions distributed across a single adult *Opn4-Cre/GFP* retina; Figure 4—figure supplement 1C). Therefore, whereas Tbr2 is expressed in a broad range of types that includes the entire ipRGC family, Tbx20 expression is confined to a diverse subset of ipRGC subtypes.

### Molecular diversity of Rasgrp1 and Tbx20 expression in ipRGCs

Our expression studies revealed that Rasgrp1 and Tbx20 have a strikingly similar pattern of expression among ipRGC subtypes. Both genes were expressed in the majority of M1 cells, a minority of M2 cells, and a small population of M5/6 cells, but not in M4 cells (Figures 4 and 5). To directly test for co-expression, we compared and contrasted the expression patterns of Tbx20- and Rasgrp1-immunoreactivity in the context of the M1-M4 subtypes revealed by Opn4-immunoreactivity (Figure 5). Rasgrp1 co-expression with Tbx20 was only observed in a fraction of M1/3 cells (26.0±1.8%; *n* = 241 across two retinas; Figure 5). Further, M1 cells expressing either Rasgrp1 or Tbx20 alone accounted for roughly similar fractions of M1 cells (37.3±4.0% and 31.1±3.5%, respectively). Only a small fraction of M1 cells were immunonegative for both Rasgrp1 and Tbx20 (5.3±0.7%). In contrast, half of M2 cells lacked Rasgrp1 and Tbx20 immunoreactivity (57.7±3.3%; *n* = 388 across two retinas). Approximately a third of M2 cells expressed Tbx20 (31.0±0.7%), while only 11.3±5.1% expressed Rasgrp1. We did not observe any example of an M2 cell expressing both Rasgrp1 and Tbx20.

**Figure 5.**
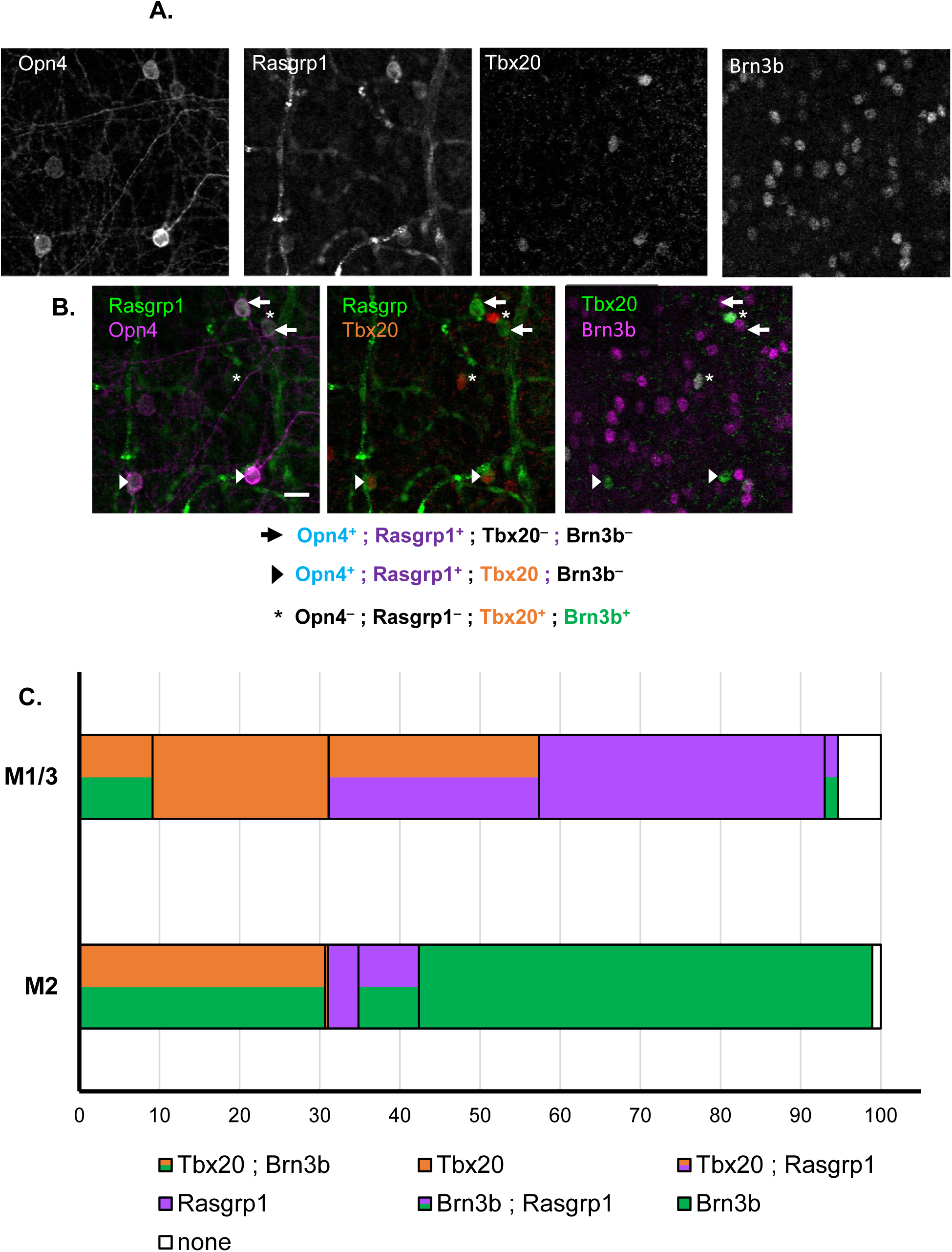
Complex pattern of Rasgrp1-Tbx20-Brn3b co-expression suggests further diversity in ipRGC family. A. Quadruple immunofluorescence study of Tbx20, Brn3b, Opn4, and Rasgrp1 (gray-scale). B. Rasgrp1 and Opn4 (left panel) were initially quantified for ipRGC subtype expression prior to comparison with Tbx20 (middle panel) and Brn3b (right panel) expression. Rasgrp1, Brn3b, and Tbx20 expression are partially overlapping. C. Integrated co-expression patterns of Brn3b, Rasgrp1, and Tbx20 with M1 and M2 ipRGC subtypes. The M1 group includes displaced M1 and M3 types.

### Molecular diversity of SCN-projecting ipRGCs

We further examined the Rasgrp1- and Tbx20-expressing ipRGC subtypes to seek intersectional expression patterns that would divide ipRGCs by their downstream visual pathways. Earlier studies showed that M1 cells could be subdivided based on their level of expression of Brn3b (Chen et al., 2011). We used quadruple immunolabeling to simultaneously test Brn3b expression with Rasgrp1- and Tbx20-immunoreactivity in the context of Opn4-immunolabeled ipRGCs (25 regions, three wild type retinas; Figure 5A,B). We determined that a minority of M1/3 cells express Brn3b (7.9±6.0%, n=241), which is similar to previous studies (Jain et al., 2012). The Brn3b^+^ M1/3 cells expressed either Tbx20 or Rasgrp1 (91.0 and 9.0±10.1%, respectively; n=30) (Figure 5C).

Further, we determined that most M2 cells expressed Brn3b (90.8±6.9, n=168). In contrast to M1/3 ipRGCs, the majority of Brn3b^+^ M2 ipRGCs did not express either Rasgrp1 or Tbx20 (67.4.6±13.8, n=222). Most M2 cells expressing Tbx20 were also Brn3b immunopositive (84.5±25.1, n=118). The small subset of M2 cells that express Rasgrp1 could be further divided by Brn3b presence or absence (5.0±6.0% and 4.5±1.4%, respectively; n=168). Generally, we found no cells co-expressing all three genes (n=729).

Our immunostaining study in the retina suggested that SCN-projecting M1 cells (Brn3b-negative) might represent molecularly diverse cell populations (Figure 5C). We correlated gene expression of Rasgrp1 and Tbx20 in the retina with retrograde labeling from the SCN (Figure 6A). We injected rhodamine-conjugated retrobeads in the SCN, followed by immunofluorescence labeling for Opn4, Rasgrp1, and Tbx20 (Figure 6A-C). All injection sites clearly involved the SCN, as revealed by DAPI labeling, but did not spread to the optic chiasm or tract (Figure 6B). Quantitative co-expression analysis (18 confocal images collected across the contralateral and ipsilateral retinas) revealed that nearly all retrolabeled cells were Opn4-immunopositive (95%). These cells exhibited variable patterns of staining for the other proteins. Most cells expressed both Rasgrp1 and Tbx20 (76%), but equal minorities expressed either Rasgrp1 (12%) or Tbx20 (12%, Figure 6D). This expression pattern was consistent across the ipsi- and contralateral retina (Figure 6D), as suggested by a bilateral input to the SCN (Hattar et al., 2006; Fernandez et al., 2016). Therefore, we show that SCN-projecting ipRGCs have a complex pattern of Rasgrp1 and Tbx20 gene expression. Together, these results provide evidence for previously unrecognized molecular diversity in adult ipRGCs.

**Figure 6.**
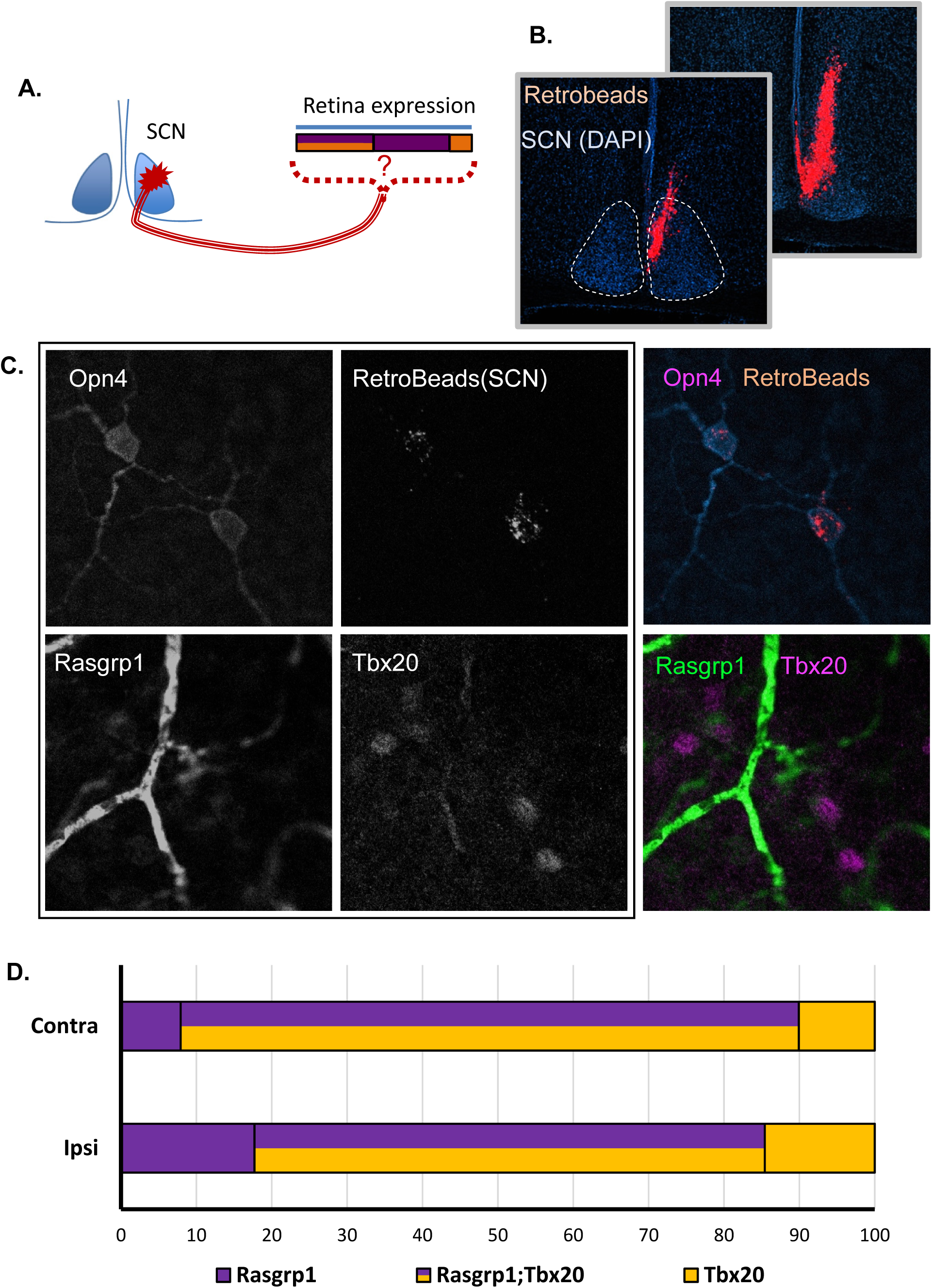
The ipRGCs projecting to the suprachiasmatic nucleus (SCN) have a molecularly diverse pattern of Rasgrp1 and Tbx20 expression. A. Experimental design of fluorescent bead injection to SCN, followed by examination of Rasgrp1 and Tbx20 expression in retrograde labeled RGCs. B. Neuroannatomical study to verify that retrograde injection is within the SCN, but not the optic nerve. B. Triple immunofluorescence of Opn4, Rasgrp1, and Tbx20 in combination with fluorescent Retrobeads. Retrobeads were mostly observed in Opn4-immunopositive RGCs (M1-M3 ipRGCs). Quantification of Rasgrp1 and Tbx20 in retrolabeled cells. D. SCN-projecting ipRGCs in the ipsilateral and contralateral retina are molecularly diverse for Tbx20 and Rasgrp1 expression.

## Discussion

Prior efforts to assess the distinctive genetic composition of ipRGCs have been complicated by their rarity among diverse retinal cell types and the inherent difficulties of maintaining viability of dissociated mature neurons (Lobo et al., 2006). Our approach was first to isolate RGCs by immunoaffinity, then to further purify ipRGCs from these based on genetic labeling and FACS, and to finally to compare the transcriptional profiles of the purified ipRGCs to those of the residual cell pool, consisting mainly of conventional RGCs. The relative purity of our ipRGC sample is supported by enrichment for transcripts of genes known to be differentially expressed in ipRGCs and the low levels of transcripts selectively expressed in potentially contaminating populations, including the abundant rod photoreceptors. Our isolation method and differential expression analysis allowed us to identify more than 75 differentially expressed genes in ipRGCs relative to conventional RGCs.

### Genes differentially expressed in adult ipRGCs

There is limited knowledge of specific gene expression in ipRGCs generally and within particular ipRGC subtypes, especially non-M1 ipRGCs. Many diverse genes appeared more highly expressed in ipRGCs than in conventional RGCs. We confirmed differential protein expression in ipRGCs immunohistochemically for two of these genes: Tbx20, a transcription factor implicated in visual development; and Rasgrp1, a G-protein exchange factor that may interact with the melanopsin phototransduction cascade. However, only a subset of ipRGCs appeared to express detectable levels of these proteins, and such variable expression was apparent even among ipRGCs of the same subtype. Some ipRGCs expressed both proteins, but many did not. This diversity even extended to the M1 cells projecting to the SCN, which had been thought to share the distinctive molecular feature of little or no expression of the transcription factor Brn3b. These novel markers of molecularly distinctive ipRGC varieties open the way for cell-type-specific manipulations through intersectional strategies.

### What type(s) of adult ipRGCs express Rasgrp1?

Rasgrp1 expression has previously been detected in the hippocampus, striatum and olfactory regions of the brain (Ebinu et al., 1998; Toki et al., 2001), but our study appears to be the first to explore Rasgrp1 expression in the eye. Rasgrp1-like immunoreactivity marked a diverse subpopulation of ipRGC subtypes, including the M1-M3 ipRGC subtypes but not the M4-type. Either M5 or M6 ipRGCs, or both, also appear likely to express Rasgrp1, because some Rasgrp1 immunoreactive cells had weak Opn4-immunoreactivity without the characteristic dendritic labeling of M1-M3 ipRGCs and with somas too small to be M4 cells (Ecker et al., 2010; Quattrochi et al., 2018; Stabio et al., 2018).

### What is the function of Rasgrp1 in adult ipRGCs?

The function of Rasgrp1 in ipRGCs is unknown, but it could interact with the melanopsin phototransduction cascade. The direct photoresponse of ipRGCs appears to increase levels of both DAG and calcium. Both of these signaling molecules bind to and activate Rasgrp1, and trigger its translocation to the Golgi apparatus (Bivona et al., 2003; Graham et al., 2008; Zhang et al., 2010). However, ipRGC phototransduction Rasgrp1 signaling does not appear to be essential for ipRGC phototransduction because more than a quarter of M1 ipRGCs and the great majority of M2 cells are immunonegative for Rasgrp1. Even in ipRGCs, Rasgrp1 may be activated by DAG and calcium signals unrelated to Opn4 phototransduction, and such signals are presumably also responsible for modulating Rasgrp1 in cells (such as certain amacrine cells) which express Rasgrp1 but not melanopsin.

Rasgrp1 has the potential to affect any number of neuronal signaling pathways. Ras signaling pathways are enormously complex and the cross talk between pathways makes it even harder to identify specific effects. One basic mechanism for specificity in Ras signaling is the distinct subcellular targeting of downstream components of the signaling pathway. In lymphocytes, localized Ras signaling of Rasgrp1 occurs preferentially on the Golgi apparatus, which is a rare form of compartmentalized Ras signaling (Bivona et al., 2003; Zhang et al., 2010). The Golgi apparatus in neurons provides the posttranslational protein modifications required for organizing protein and organelle trafficking throughout the cell. Rasgrp1 could play a crucial role in orchestrating a specific set of post-translational modifications at the Golgi.

An important survival mechanism in M1 ipRGCs is the maintenance of mTOR expression by melanopsin phototransduction (Duan et al., 2015; Li et al., 2016). The DAG-activated Rasgrp1-Ras-Mek1/2-Erk1/2 pathway contributes to mTOR activation in thymocytes (Gorentla et al., 2011). The percentage of M1 cells expressing Rasgrp1 (72%) in this study is similar to the percentage of M1 cells that survive optic nerve crush (∼70%, Li et al., 2016). It would be interesting to test whether Rasgrp1 is required for the maintained mTOR levels in these surviving M1 ipRGCs. Recent studies have also clarified different mechanisms responsible for M1 survival and the regeneration potential of alpha-RGCs such as M4 ipRGCs (Duan et al., 2015; Li et al., 2016). Despite surviving well, M1 ipRGCs were not capable of regenerate their axons (Li et al., 2016). In contrast, alpha-RGCs had a unique capability to regenerate their axons that was promoted by their high levels of mTOR expression as well as osteopontin/Spp1 expression coupled with the growth factor IGF-1(Duan et al., 2015). Whereas M1 ipRGCs were found to maintain mTOR expression after axon injury, M4 ipRGCs and other alpha RGC types were demonstrated to have diminished mTOR expression levels (Li et al., 2016). The absence of detectable Rasgrp1 expression in M4 ipRGCs may help to account for their failure to maintain mTOR expression after injury.

### Central brain targets of Rasgrp1-expressing ipRGCs

At the circuit level, Rasgrp1 is not anticipated to be an essential regulator of circadian photoentrainment. The majority of M1 ipRGCs are known to provide the primary retinal input to the SCN, while a subset of M1 ipRGCs send input to the OPN to regulate the pupillary reflex. Our studies found that Rasgrp1 was expressed in the majority of M1 ipRGCs, which suggested to us that it might correlate directly and completely with the SCN-projecting M1s. However, retrograde tracing experiments from the SCN revealed that some SCN-projecting M1 cells were Rasgrp1-immunonegative. Accumulating research suggests that the SCN is more compartmentalized than previously recognized (Bedont and Blackshaw, 2015). The question of whether Rasgrp1 is expressed in a subset of M1 ipRGCs that targets a specific compartment of the SCN remains to be determined.

We find Rasgrp1 to be expressed not only in SCN-projecting M1 ipRGCs, but also in other ipRGC subtypes, especially M2 cells and apparently M5 and/or M6 cells. Collectively, these types project to various non-image-forming visual centers, including the olivary pretectal nucleus, intergeniculate leaflet, and dorsal lateral geniculate nucleus (Quattrochi et al., 2018; Stabio et al., 2018).

### Tbx20 is expressed in a diverse set of ipRGC subtypes

The T-box transcription factor Tbx20 exhibited enriched expression relative to conventional RGCs in postnatal and adult retinas. Double immunolabeling revealed that many ganglion cells that expressed this protein also expressed melanopsin. As was true for Rasgrp1, Tbx20 was determined to be expressed in most M1 ipRGCs (75%), a substantial minority of M2 cells (40%) and an additional population of RGCs whose identity was not immediately obvious. We decided to then compare Tbx20 against other known molecular patterns in ipRGCs. Whereas most RGCs follow a Brn3b-dependent developmental program, the M1 ipRGCs that project to the SCN do not express Brn3b while OPN-projecting M1 ipRGCs express Brn3b. We found that Brn3b-expressing M1 cells are also Tbx20-immunopositive. The Brn3b-negative M1 cells are split between cells that express Tbx20 and those that do not. This finding suggests that ipRGCs are more molecularly diverse than originally anticipated: Tbx20 is expressed in ipRGCs with differing brain targets, Tbx20 is expressed across multiple morphologically defined subtypes, and Tbx20 is not expressed in all of any one type. The exploration of Tbx20 coexpression with Rasgrp1 revealed a complex coexpression pattern among M1-M3 ipRGCs.

### What is the function of Tbx20 in adult ipRGCs?

Tbx20 has well-established roles in embryonic development and is continuously required in mature neurons and other cell types to maintain their identities and functional properties during adulthood (Naiche et al., 2005). Tbx20 functions as a repressor in early embryonic ocular development (Carson et al., 2000; Pocock et al., 2008) and is required for the expansion of the small pool of precursor cells in the optic vesicle (Carson et al., 2004). However, little is known about the function of Tbx20 in the adult retina.

Tbx20 can function as a transcriptional activator in parallel with its repressor activity, with these two roles impinging on distinct biological processes (Sakabe et al., 2012). In addition to its key developmental roles, Tbx20 appears vital for maintaining the structure and function of cardiac muscle cells in the adult mouse heart (Stennard et al., 2003; Shen et al., 2011). In adult cardiomyocytes, Tbx20 is responsible for directly activating genes critical for normal adult cardiac function such as those required for ion transport and heart contraction (Shen et al., 2011; Sakabe et al., 2012). In contrast, genes directly repressed by Tbx20 have known roles in non-heart developmental programs, cell cycle, proliferation, and immune response (Sakabe et al., 2012). The transcriptional effects of Tbx20 shift during cardiac development, from early mediation of proliferation of cardiac progenitors, to implementation of an anti-proliferative program in the adult heart (Cai et al., 2005; Takeuchi et al., 2005). Therefore, Tbx20 cooperates with distinct cohorts of transcription factors to either promote or repress distinct molecular programs in a context-dependent manner (Sakabe et al., 2012). Similarly, binary cell fate specification in the retina is driven by complex genetic programs that require the simultaneous activation and repression of genes by transcription factors. Tbx20 may prove to have a similar reversal in its transcriptional activity in the retina when transitioning from broad embryonic development program to regulating adult neuron identity of a subset of ipRGCs. Further, Tbr2 is another Tbox family member that is known to have a critical role in the development of retinal ganglion cells (Mao et al., 2008), which later becomes essential to a restricted set of ipRGCs that participate in NIF visual circuits (Mao et al., 2014; Sweeney et al., 2014). Our studies determined that Tbx20 and Tbr2 are coexpressed in adult ipRGCs. They may work cooperatively to specify ipRGC subtype identity by regulating cell-specific transcriptional programs and repressing alternate fates.

### Characterization of ipRGC subtypes

Retinal cell types are generally classified using a combination of morphology, gene expression, mosaic organization, light responses and synaptic connectivity (Sanes and Masland, 2015). By these criteria, intrinsically photosensitive RGCs comprise at least 6 distinct cell types (Figure 7). Though all express melanopsin, they differ from one another in the strength of the intrinsic response, their morphology, pattern of light responses, and projections to the brain. However, the further subdivision may be in order. The M1 type appears subdivisible into at least two subtypes, one expressing the transcription factor Brn3b and innervating the OPN and geniculate complex, while the other lacks Brn3b expression and innervates the SCN (Chen et al., 2011). Our study shows further diversity in the M1 and M2 types based on the expression of Rasgrp1 and Tbx20. For example, we find molecular diversity among in the SCN-projecting ipRGC subtypes (Figure 7). It is unclear to us whether this should be used to propose a further formal subdivision of M1 and M2 cells. For example, the expression of these proteins could fluctuate over time in individual cells and be uncorrelated across cells of the same type. One would like to know that these patterns of expression are stable over time and correlated with other cell features before proposing such further subdivision. The extensive overlap among dendritic fields of M1 (and of M2) cells (Berson et al., 2010) means that either type could be subdivided into two or perhaps three subtypes while still maintaining full retinal coverage (i.e., tiling), but it seems likely that we will either have to accept that there is substantial molecular diversity with single types (as there is substantial functional diversity among M1 cells (Emanuel et al., 2017)) or that the dogma of complete retinal tiling by single types must be abandoned.

**Figure 7.**
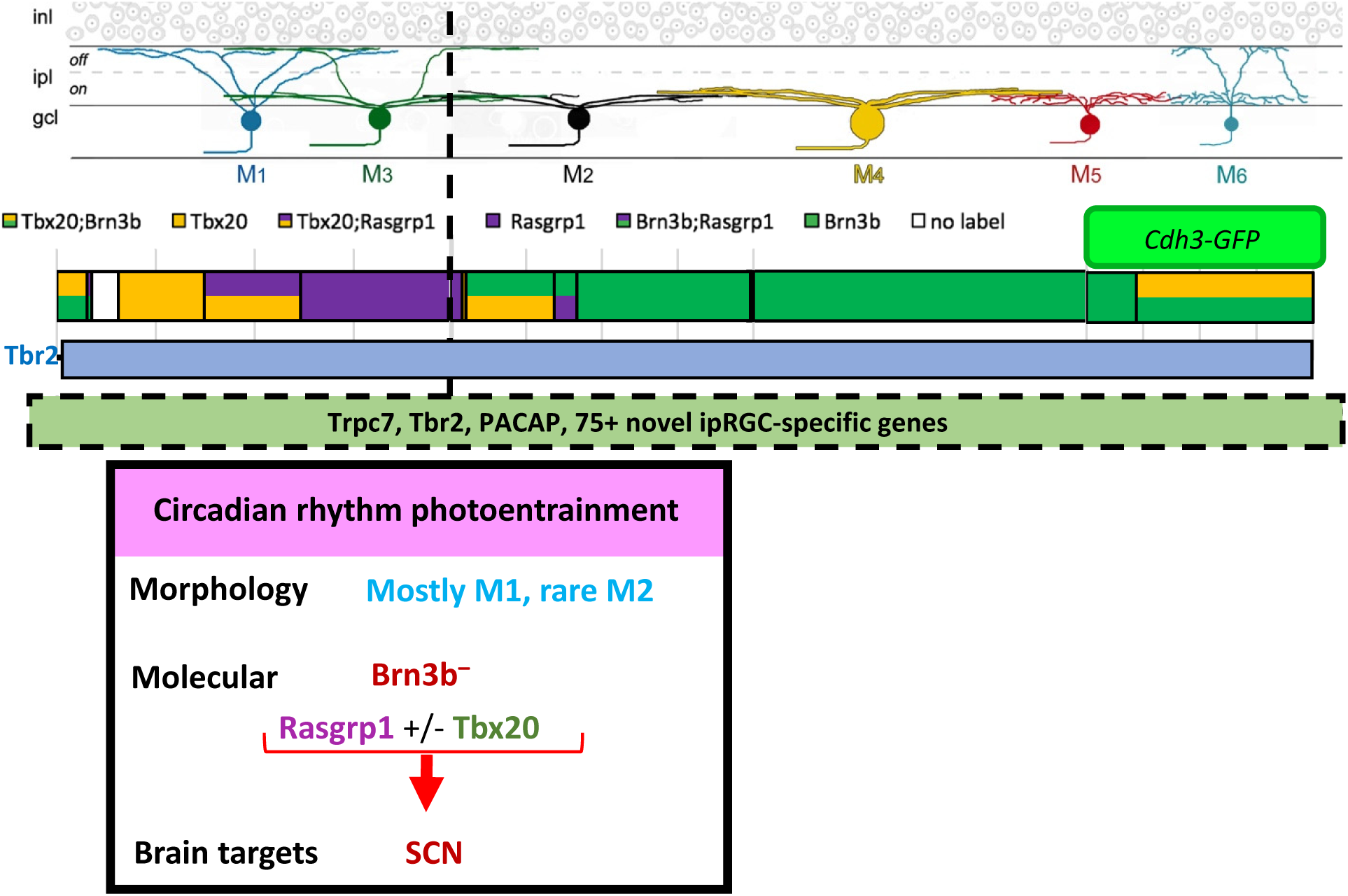
Current model of ipRGC family members integrating molecular, physiology, brain circuitry, and morphology (see text for details).

In addition to Tbx20 expression in a subset of SCN-projecting M1s, Tbx20 may also regulate the molecular program of ipRGCs projecting to the OPN and control the pupillary light reflex. The Brn3b-expressing M1 ipRGCs, a subset of M2 ipRGCs and *Cdh3-GFP* labeled M6 ipRGCs are all known to project to the OPN, and all express Tbx20. However, this is not a direct association and further retrograde or Tbx20 conditional knockout studies are required, especially to determine the function of Tbx20-expressing M2s.

### Comparison with other ipRGC gene expression profiles

Siegert and colleagues (2012) surveyed gene expression in many different sets of mouse retinal neurons, using specific mouse reporters strains (including the *Opn4-Cre* reporter system for ipRGCs), FACS isolation of labeled cells, and microarray analysis. Many of the genes they found strongly expressed in ipRGCs were also among the genes we found differentially expressed in ipRGCs. However, dozens of additional genes differentially expressed in ipRGCs emerged from our analysis that were not detected in theirs (Siegert et al., 2012). Discrepancies between their findings and ours may stem from technical factors such as differing degrees of contamination of the starting material with rod photoreceptor transcripts, the use of internal control cell populations for relative gene expression comparison in our study but not theirs, and differences between microarray and RNA-sequencing methodologies.

Another recent studied used single-cell transcriptomic analysis of the mouse retina and were able to identify ipRGCs from their cell suspensions (Macosko et al., 2015). Single cell technology is ideal, in principle, for the precise identification of an individual neuron’s molecular identity despite the extreme heterogeneity of the nervous system. Macosko et al., 2015 could definitively distinguish the main broad class of RGCs, but they required *post hoc* supervised analysis to distinguish a limited number of genes attributed to Opn4-positive cell clusters. They identified nine genes with a two-fold increase in expression compared to Opn4-negative cells. Three genes (Tbr2, Igf1, and Tbx20) were also found to be enriched in our ipRGC samples. In contrast, Tbx20 did not reach above threshold for Siegert et al., (2012), but it is among the highest expressing ipRGC-enriched genes in our study. Single-cell gene expression profiling methods such as Drop-Seq hold great promise and will certainly continue to be pursued more in future studies of neuron subtype gene expression. Recently, single-cell studies of pre-enriched bipolar cells were able to distinguish molecular markers for all previously recognized bipolar cell types (Shekhar et al., 2016). Similarly, ipRGCs subtype-specific gene expression may become deciphered in the future using single-cell analysis that incorporates a pre-enrichment step for ganglion cells.

## Conclusion

In conclusion, our results demonstrate a method to purify ipRGCs and identify an extensive list of more than 75 genes that are differentially expressed compared to generic RGCs. Of course, our identified differentially expressed genes in ipRGCs should not be considered a complete account of all genes relevant to ipRGC function. Instead, we hope that it will provide a beneficial resource that will generate hypothesis that lead to key insights into ipRGC function in non-image forming vision. The more than 75 genes suggested to be differentially expressed in ipRGCs will be useful for the identification of marker genes for ipRGC subtypes, comparison of gene expression across types, understanding the intracellular gene networks underlying ipRGC phenotypes, and the testing for conservation of ipRGC molecular programs across mammalian species.

We are encouraged that our gene expression profiling data of ipRGCs lead us to the identification of Tbx20 and Rasgrp1 as novel, selectively expressed genes in ipRGCs. These results serve as a good proof of principal for the validity of our gene expression profiling results. In addition, the stable, specific expression in adult ipRGCs suggests that the differential gene expression of Rasgrp1 and Tbx20 were not simply due to transient, stress-induced molecular program resulting from the harsh processing steps prior to sequencing. We determined that Tbx20 and Rasgrp1 are expressed across ipRGCs that belong to multiple morphological types, have diverse molecular expression, differing physiology, and are involved with multiple visual brain circuits.

## Methods

### Animals

All experiments were conducted in accordance with NIH guidelines under protocols approved by the Brown University (Providence, RI) Animal Care and Use Committee. Both male and female adult mice (P30 to P90) were used unless otherwise stated. *Opn4*^*cre/cre*^ mice (Ecker et al. 2010) crossed with floxed-stop reporter mice: “*Z/EG*” (*Jax#003920*); the offspring express GFP in *cre*-expressing cells (M1-M6), as described by Ecker et al., 2010. *Opn4-GFP(ND100Gsat)* is a BAC transgenic originated from the GENSAT project at Rockefeller University. Rasgrp1-KO (Rasgrp1^tm^1^Jstn^, Dower et al., 2000) was initially provided generously by Robert Barrington, U. of Alabama for initial testing. A colony was created inhouse from stock at Jackson labs (Jax 022353). *Rasgrp1-Cre*(PO1 founder line) was rederived from Jackson Labs (#34811-UCD). *Cdh3-GFP* reporter is a BAC transgenic originally generated by the Gensat project (MMRRC, BK102Gsat/MMNC) and has been used previously (Quattrochi et al., 2018). This mouse line has been backcrossed in wild type background for at least 10 generations. *Cdh3-GFP* mice were three weeks old or younger unless otherwise stated.

### Gene expression analysis of purified mouse ipRGCs

For our transcriptomics studies, we used two ipRGC reporters available in the lab to identify selective gene expression in ipRGCs compared to RGCs as a whole: 1) BAC transgenic *Opn4-EGFP* Gensat mice from MMRRC (#033064-UCD) and 2) knock-in *Opn4-Cre* mice (obtained from S. Hattar; Ecker et al., 2010) with Cre-dependent GFP reporter (Z/EG obtained from Jackson labs; (Novak et al., 2000). The *Opn4-GFP* mice were maintained as heterozygotes on *C57/BL6* background while *Opn4*^*Cre/+*^;*Z/EG* mice were generated by breeding homozygous *Opn4-Cre* mice with heterozygous *Z/EG* mice and maintained on mixed background. Although *Opn4-GFP* reporter labeling in ipRGCs has not been reported previously, a similar BAC transgenic strategy has been demonstrated to label M1-M3 ipRGCs (Schmidt et al., 2008). We tested the correlation of the *Opn4-GFP* reporter expression and endogenous melanopsin in RGCs. We used anti-melanopsin immunoreactivity label M1-M3 ipRGCs (Berson et al., 2010) and anti-EGFP antibodies in a whole-mount retina from an adult *Opn4-EGFP* mouse. Importantly, all *Opn4-GFP*^+^ cells were Opn4-immunopositive (n=60 GFP^+^ cells across seven regions of one retina, Figure 8A). Unexpectedly, more than half of the Opn4-immunopositive M1-M3 ipRGCs were not labeled by the reporter (55%, n=133). Therefore, the coexpression of EGFP expression by the *Opn4-GFP* reporter is strongly correlated with M1-M3 ipRGCs, but only accounts for about half of the population. The other reporter system used in this study, *Opn4*^*Cre/+*^; *Z/EG* mice, has been previously demonstrated to label with EGFP all six known morphological types of ipRGCs), named M1–M6 (Ecker et al., 2010; Quattrochi et al., 2018; Stabio et al., 2018). However, Opn4-immunofluorescence studies of four regions across an *Opn4-Cre/GFP* retina revealed that more than one-fourth of M1, displaced M1, and M2 cells lacked discernable GFP-labeling (28%, n=81; 27%, n=15; 33%, n=132, respectively) (Figure 8C). There were many additional GFP^+^ cells that were Opn4-immunonegative, with large soma cells being designated as M4 ipRGCs (54%, n=67 M4 cells). The remaining identified M4 cells had somas with weak, but present, Opn4-immunoreactivity and were only partially accounted for by GFP-labeling (27% Opn4^+^;GFP^neg.^, n=67 M4 cells). Additionally, small soma GFP^+^ cells with absent Opn4-immunoreactivity were designated as M5/6 cells, since the lack of dendritic labeling made it impossible to distinguish between the M5 and M6 types (72%, n=202 M5/6 cells). We observed small cell bodies with weakly Opn4-immunlabeling that did not extend to the dendrites, which were also designated as M5/6 types (28%).

**Figure 8.**
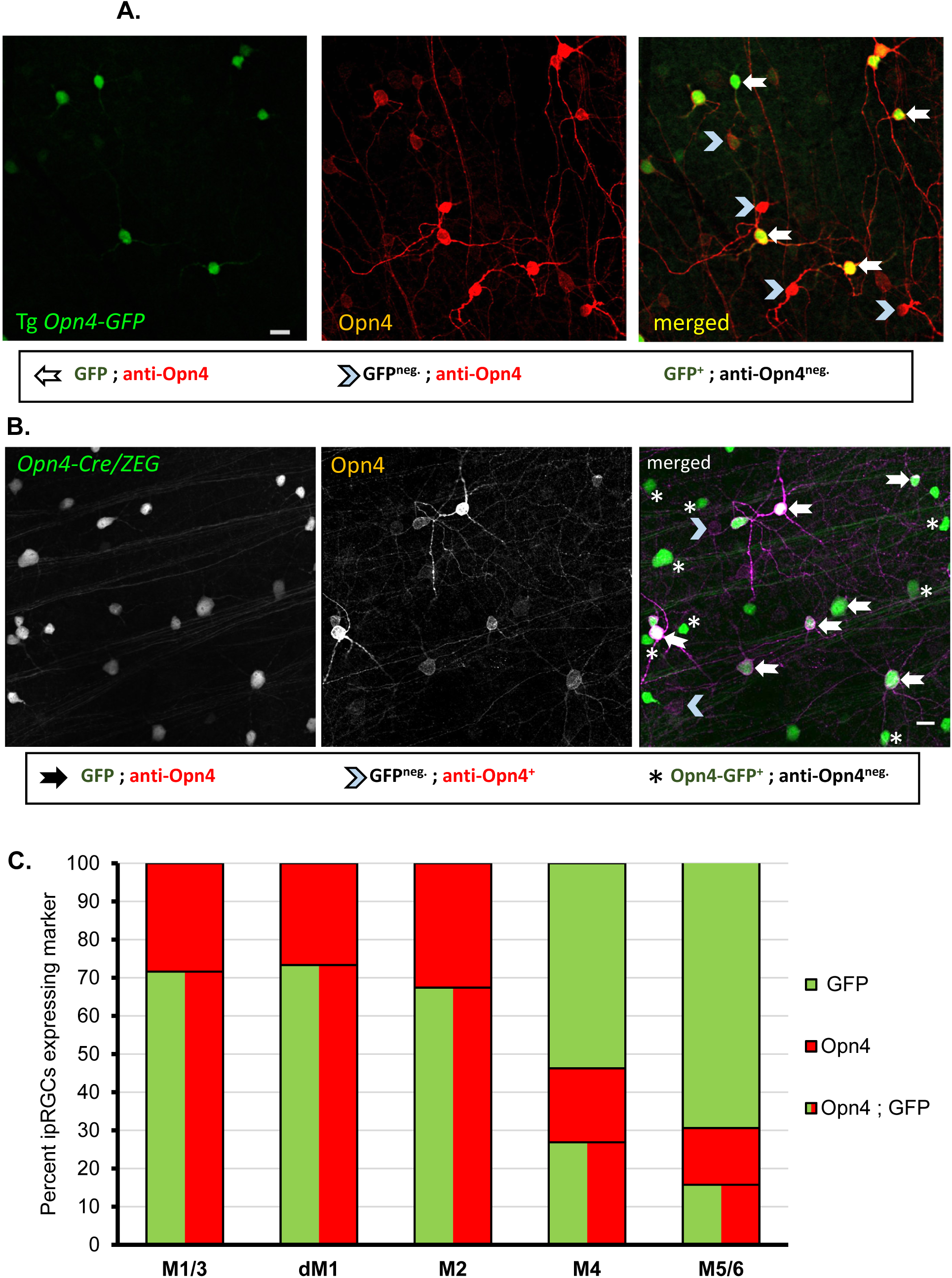
Coexpression of BAC transgenic Opn4-GFP reporter with Opn4 immunoreactivity. A.Immunofluorescence of anti-Opn4 staining of whole mount retina from transgenic *Opn4-GFP* mice with fluorescent protein expression in ipRGCs. Red, Opn4-immunolabeling; green, fluorescently labeled cells; yellow, merged co-localized labeling pattern. Scale bar, 20 μm. B. Co-expression of *Opn4-Cre/GFP* labeling with immunofluorescence of anti-Opn4 staining of whole mount retina. Red, Opn4-immunolabeling; green, fluorescently labeled cells; yellow, merged co-localized labeling pattern. Scale bar, 20 μm. C. Quantification of labeling efficiency of Opn4-immunolabeled M1-M3 ipRGCs by *Opn4-Cre/GFP*. Additional comparison of GFP-labeling in low Opn4-expressing ipRGC subtypes M4 (large soma) and M5/6 (small soma).

Two ages were chosen for retina tissue collection for purification of ipRGC neurons: Postnatal day 5 (+/- 1day) and young adult (P30 +/- 3 days). Three or more biological replicates were used for each dataset (three for postnatal day 4-6 (P4-6) Gensat *Opn4::GFP*, six replicates for Gensat *Opn4::GFP*, four replicates for *Opn4::Cre/GFP* reporter). Each adult replicate required the pooled retinas from 15-20 transgenic mice to acquire suitable numbers of cells for the transcriptomics. These steps are outlined in much more depth and detail in the following sections.

The choice of transcriptomics preparation strategies and final readout of processed samples are interrelated and subject to a number of technical concerns vital to the success of transcriptomics analysis. Seven steps in the development and completion of the cell type-specific transcriptomics procedure of cells isolated from early postnatal and adult mouse retinas (Figure 1). **First**, dissociation of retinal tissue to a cell suspension; **Second**, purification of RGCs; **Third**, FACS analysis and sorting; **Fourth**, extraction of RNA; **Fifth,** cDNA amplification because the resulting RNA is typically low in abundance; **Sixth**, shear cDNA into sequenceable fragments that are then sequenced with Illumina deep-sequencing; and **Finally**, the raw reads must be analyzed to identify differential expression of genes in ipRGCs.

### Retina dissociation

The intertwined nature and tight cell–cell adhesions of neural cells make it difficult to separate cells without causing cellular damage and activating stress or cell death pathways. Therefore, vigorous dissociation of tissues can lead to activation of stress or cell death pathways and distort the resulting expression profile. However, poor dissociation can lead to a severe decrease in isolated cells and make downstream expression studies an impossibility. As a first step, we dissected retinal tissue, which we then digested in a protease solution to loosen and disrupt the intertwined neural cells into single cell suspensions for subsequent cell-type isolation. The essential components for proper dissociation included: the proteolytic enzyme papain L-cysteine to promote enzymatic activity, DNase for destroying the extremely sticky free DNA strands released by damaged cells, and an absence of calcium from solutions to promote disruption of cell-cell adhesions (Barres et al., 1988). We further optimized cell viability and recovery by replacing dPBS with HibernateA and including B27 throughout the cell-isolation procedure (Brewer et al., 1993; Brewer, 1997; Brewer and Torricelli, 2007). The use of survival-promoting media such as Hibernate-A and supplements such as B27 was particularly relevant for adult neural dissociation since adult neurons have been demonstrated to be especially prone to cell death (Eide et al., 2005; Brewer and Torricelli, 2007). We incubated the dissected retinas in pre-activated protease solution and completed the cell dissociation by triturating the cells with a 1mL pipette tip.

### RGC Pre-Enrichment

Pre-enrichment of ganglion cells prior to isolation of ipRGCs is necessary due to their extraordinary rarity in the retina. Ganglion cells only make up 1% of mouse retinal cells and only 1-5% of ganglion cells are ipRGCs (0.01-0.05% of retina cells) (Hattar et al., 2002; Ecker et al., 2010; Berson et al., 2010). In contrast, the classic photoreceptors rods and cones account for 75-80% of all mouse retinal cells (Jeon et al., 1998). Therefore, we pre-enriched for ganglion cells prior to positively selecting fluorescently labeled ipRGCs using fluorescence activated cell sorting. We incorporated an immunopanning system that has proven effective at isolating a homogeneous population of RGCs (Barres et al., 1988; Cahoy et al., 2008). Immunopanning takes advantage of the cell surface protein Thy1 which is shared among retinal ganglion cells (Barres et al., 1988). The classic immunopanning process uses antibodies raised against Thy1 to select the RGCs from the heterogeneous retinal cell suspension. We improved upon viability and reproducibility of the immunopanning procedure by adapting it to use a magnetic-activated cell sorting procedure (MACS, Miltenyi Biotec), eliminating the need for using harsh lysis treatment (data not shown). We incubated the dissociated retinal cell solution with Thy1.2-conjugated magnetic nanoparticles which are retained in a magnetized column and the isolated RGCs can then be acquired (Haeryfar and Hoskin, 2004). In preparation for FACS, the DNA intercalating dye, 7-AAD, was added to the solution to discriminate dying cells that consequently had compromised cell membranes.

### Fluoresence-Activated Cell Sorting (FACS)

We used a FACS Aria (BD Biosciences) electrostatic sorter to isolate a homogeneous GFP-labeled cell population from the dissociated RGC-enriched cell suspension of the transgenic melanopsin reporter mice (Figure 1). FACS has been successfully used previously to profile gene expression in cell subtypes of the nervous system (Lobo et al., 2006; Cahoy et al., 2008; Siegert et al., 2012). Live cell gating was achieved by excluding 7-AAD labeled cells (high fluorescence signal in far red emission) (Figure 9). Fluorescently labeled ipRGCs were positively selected from the solution of enriched ganglion cells by their relatively high level of GFP fluorescence (FITC gating). In parallel, cells with relatively low GFP-fluorescence (low FITC) were also isolated and designated as a “generic RGC” control population for direct comparison with correlated ipRGC samples. The parallel isolation of generic RGCs was designed as an internal negative control for comparison with isolated ipRGCs. Accordingly, the generic RGC populations were treated with the same reagents, cytometer settings, centrifugation forces, and temperatures throughout the procedure. This is especially important for isolating adult RGC populations, since they are particularly susceptible to the stresses of FACS sorting(Lobo et al., 2006; Cahoy et al., 2008; Heiman et al., 2008). The large amount of small cellular debris generated during the dissociation process made it a challenge to keep the number of collected generic RGCs consistent with the rare ipRGCs (Figure 9). Cellular debris registers as being essentially non-fluorescent, is smaller and less complex than the generic RGC population that is of the same relative size and complexity as the isolated GFP-positive cells. Therefore, we limited cell debris by acquiring the ipRGC (GFP+) and generic RGC (GFP-negative) populations with the same relative cell complexity (indicator of cell health) and cell size selection to exclude the relatively small cellular debris or doublets.

**Figure 9.**
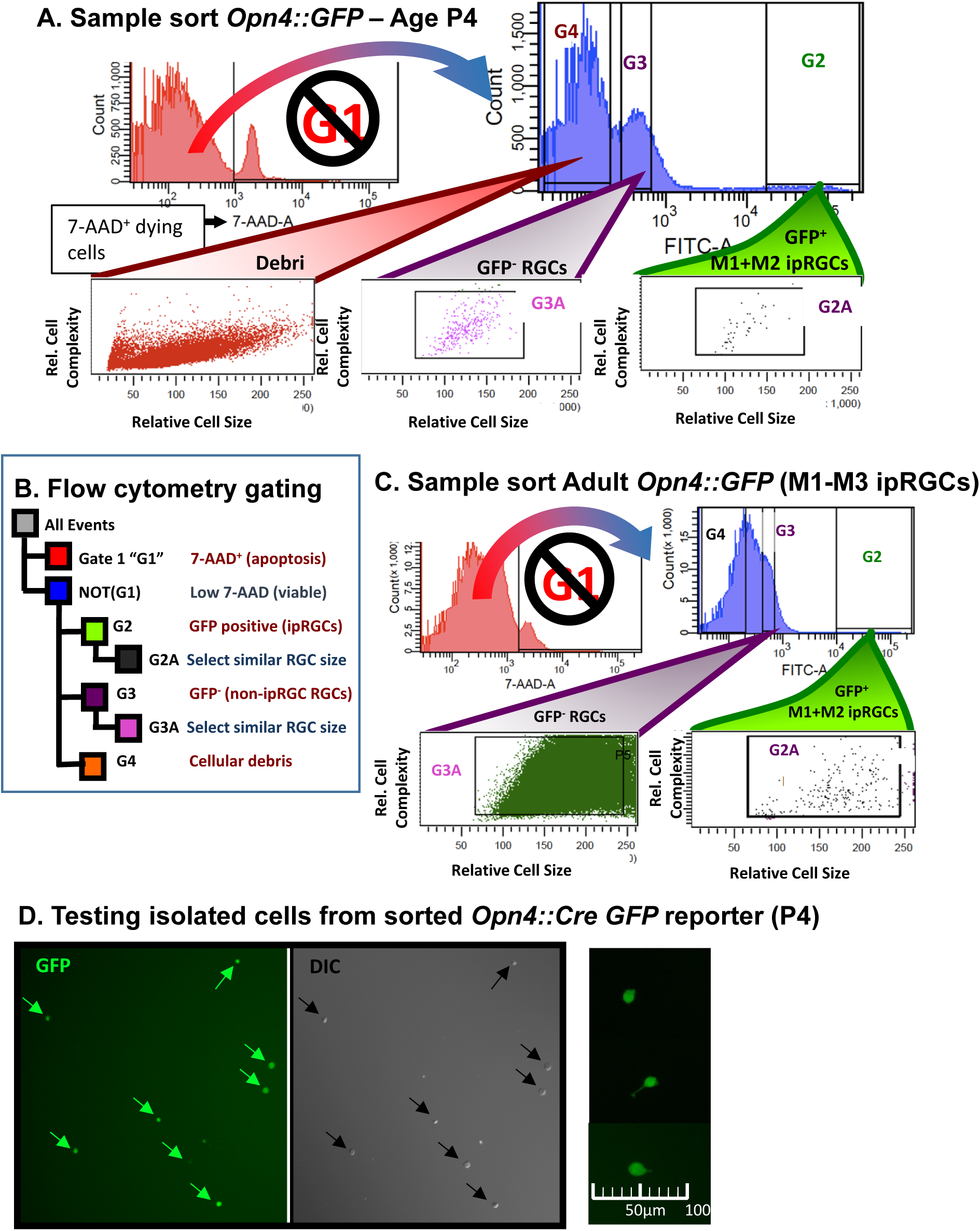
Fluorescence activated cell sorting (FACS) gating strategy for isolating ipRGCs (GFP+) in parallel with GFP-negative cells that are enriched for RGCs. A-C. Healthy cells were selected against death marker 7-AAD (not G1). The ipRGCs (GFP^+^) and generic RGCs (GFP^−^) cells were selected based on intensity and similar relative cell size ultimately using gates G2A and G3A, respectively. A. Example sort from retina of postnatal day 4 (P4) *Opn4-GFP* mouse. C. Example sort from retina of young adult *Opn4-GFP* mouse, with noticeably higher debri and cell death. D. Microscopy testing of accurate sorting of GFP+ cells isolated from P4 *Opn4-Cre/GFP* mouse.

### RNA Extraction

The small volumes allowed by the electrostatic FACSAria sorter allowed us to lyse sorted cells directly into Qiagen RLT buffer and directly proceed to RNA extraction using Minelute columns (Qiagen). The enriched RNA was treated on-column with DNase to remove any residual genomic DNA from the sample. RNA-processing was done in an enclosed RNase-free environment to limit degradation of RNA throughout the extraction process. Additionally, RNA integrity was analyzed using the Agilent 2100 Bioanalyzer and the PicoChip, which is able to qualitatively test the low RNA recovery samples (Figure 10A). Initially, we proceeded immediately with cDNA processing after RNA extraction. However, freezing at −80 degrees did not seem to effect RNA integrity since the frozen RNA samples still received RIN score of 9.0 or greater (Figure 10A). Therefore, most of the cDNA libraries were prepared after storage of extracted RNA at −80 degrees Celsius.

**Figure 10.**
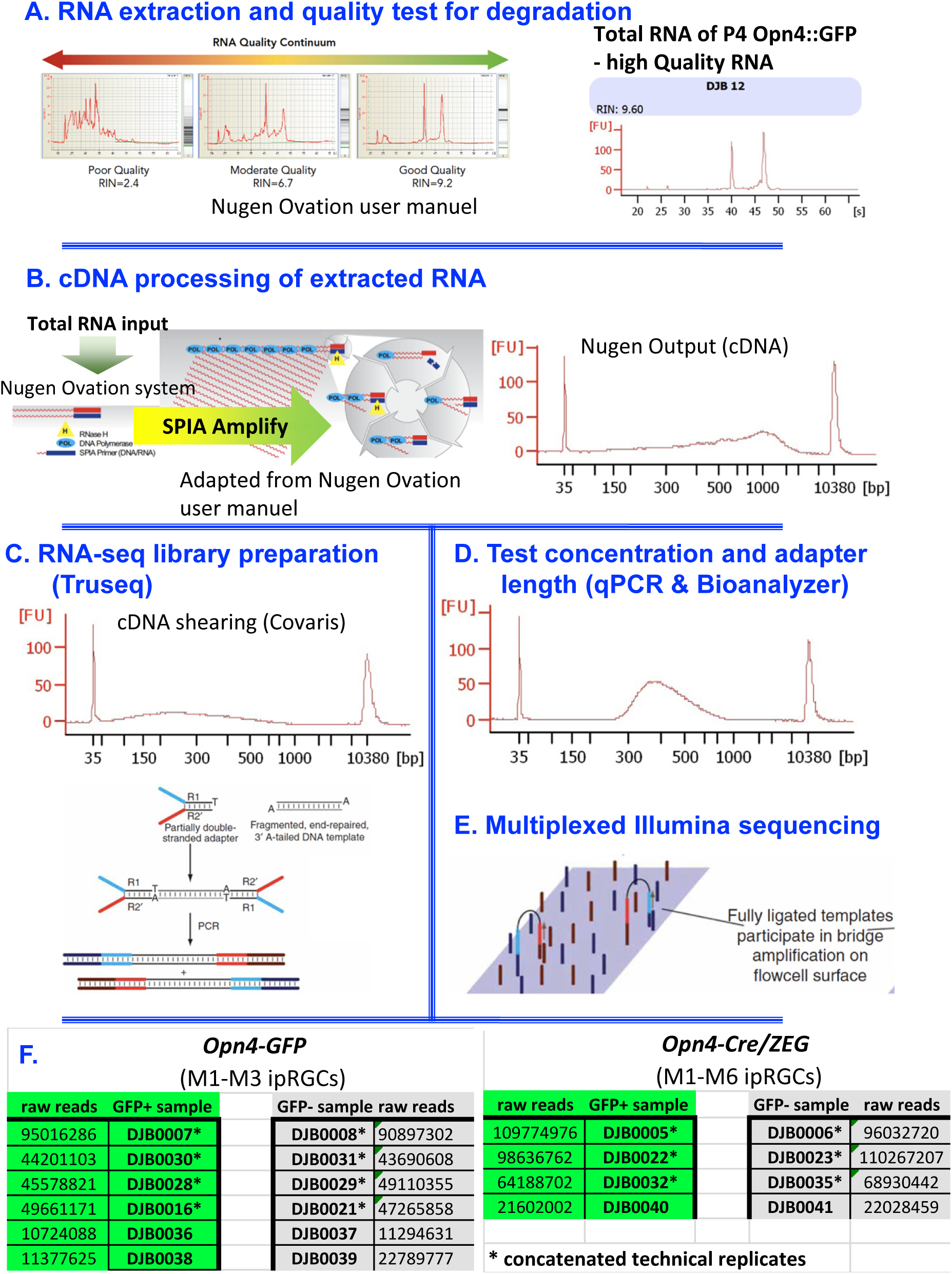
Steps involved with processing mRNA extracted from purified cell populations and preparing for RNA-sequencing (see Methods for details).

### cDNA Preparation

RNA-seq transcriptome analysis requires large amounts of RNA material using TruSeq, ranging on the order of 100-1000ng of total RNA. However, our improved method for isolating ipRGCs was able to isolate 12,000 GFP+ ipRGCs from nine postnatal day 5 (P5) transgenic reporter mice. This was only expected to provide about 12ng of extracted total RNA by qualitative estimates considering that a single cell holds 5-10pg of total RNA. Further, twice as many adult mice of the same genotype were required to provide only 1,000 cells due the the relative fragility of adult retinal ganglion cells described above. Therefore, we decided that some form of amplification was necessary to study the molecular programs used by adult ipRGCs. The Nugen Ovation RNA amplification system was successfully used in microarray studies with as little as 500pg of total RNA input (Caretti et al., 2008; Clément-Ziza et al., 2009; Morse et al., 2010) (Figure 10B). Sequencing analysis using the Ovation system has previously been reported to generate cDNA containing negligible rRNA reads (<4%), while providing a representative transcriptome with sufficient biological replicates(Tariq et al., 2011).

### RNA-seq library preparation

We determined that the Truseq system was the necessary platform for my cDNA samples to produce the 10nM sequencing library concentration required at the on-site Genomics Core facility in preparation for 50bp single-end Illumina sequencing. Before preparing the library, we first sheared cDNA to the appropriate size, (200-300bp median) using the Covaris system (Figure 10C). Each sample was subsequently processed using a unique barcode adapter to allow for multiplexing multiple samples on the HiSeq (commonly 200million 50bp reads divided among sequencing samples). Excess adapter sequences was removed using Ampure bead isolation, which removes all DNA fragments less than 200bp. Finally, the Genomics Core completed the final quality control of the DNA library prior to sequencing: testing the library fragment size distribution (High Sensitivity Bioanalyzer) and qPCR analysis using primers that match the library adapters (Quail et al., 2008) (Figure 10D).

Initially, we processed a large number of samples at once with multiplexed sequencing using HiSeq (8 samples per lane) to minimize high costs of sequencing at the expense of sequencing depth. We later reran many of the sequencing libraries with less multiplexing, enabling increased sequencing depth. The corresponding technical replicates were merged together for differential expression analysis (Figure 10E). The final read counts of each sample is shown in Figure 10F.

### Differential gene and transcript expression analysis

The completion of sequencing generated tens of millions of reads that are used to compare gene expression levels between isolated ipRGCs (GFP+) and generic RGCs (GFP-). Well-established, powerful RNA-seq differential expression analysis pipelines have been developed such as Cuffdiff, EdgeR, and DESeq (Anders and Huber, 2010; Trapnell et al., 2012; Anders et al., 2013). The Cuffdiff pipeline prioritizes isoform quantification and diversity (Trapnell et al., 2012). However, the short 50bp single-end reads that are generated in our study are not well-suited for prioritizing isoform discovery and analysis. Further, Cuffdiff does not fully take advantage of our purposeful pairwise-comparison between groups. In contrast, the EdgeR package is better suited for our purpose; it is designed to count the number of reads that align to an annotated gene (mouse reference genome, in our case) and subsequently performs statistical analysis on a generated table of counts to identify quantitative changes in expression levels between the two experimental samples. EdgeR compares and retains the relationship between all pairs of experimental samples when calculating differential expression likelihood (Anders et al., 2013). Our analysis filtered out genes with very low counts, less than 1 count per million (cpm), in more than half of the samples used in the differential expression analysis. This is a common cutoff and considers 1) that a gene must be expressed at a minimum threshold in order to become biologically important and 2) that the inclusion of genes with very low counts may negatively affect the statistical approximations used by the EdgeR pipeline.

To identify the set of differentially expressed genes in the ipRGC populations, we used the following strict criteria. First, we identified genes with low false discovery rate (FDR < 0.05) and high fold-change (greater than 2-fold) suggesting differential expression between ipRGCs and generic RGCs. Second, we considered whether the differentially gene expression was corroborated across both reporters (*Opn4::GFP* labeling M1-M3 cells and the *Opn4::Cre/GFP* system that labels M1-M6 ipRGCs) and both ages (P5 and adult). Third, we identified whether the differentially expressed genes have nearly absent gene expression in generic RGC samples to distinguish potential for selective gene expression in ipRGCs. This was distinguished both at the level of count-values and manual inspection of aligned raw reads using the Integrated Genome Viewer (IGV) (Thorvaldsdóttir and Robinson, 2013). Using IGV, we verified that the reads align with reference gene model for full-length coverage across multiple ipRGC replicates and that there were absent or partial reads aligned across the generic RGC replicates.

Determining differentially *repressed* genes in adult ipRGCs was confounded by the high amount of contaminants in generic RGC populations. We could not decipher whether a gene with relatively low expression in ipRGCs was the result of non-RGC populations contaminating the generic RGC control population. In contrast, the P5 generic RGC samples from the *Opn4-GFP* reporter were determined to have greatly reduced levels of contamination and similar levels of RGC marker expression (Figure 1—figure supplement 1). This made it possible to identify genes more weakly expressed in ipRGCs than in generic RGCs in early postnatal development (Figure 2—figure supplement 1).

### Count-based differential expression pipeline for RNA-seq data using edgeR and/or DESeq

~~~
# 1) Assess sequence quality control with FastQC)
# 2) remove adapters
$ fastx_clipper -Q33 -a adapter_sequence -l 15 -v -i DJB0005.fastq -o ad_DJB0005.fastq
# 3) remove low quality reads
$ fastq_quality_filter -v -Q33 -q 30 -p 90 -i ad_DJB0005.fastq -o adq_DJB0005.fastq #################################################################
# Align the reads (using tophat2) to the reference genome
$ tophat2 –no-coverage-search -o DJB05_th2out genome DJB05_merged.fastq #################################################################
# Sort by name, convert to SAM for htseq-count
$ samtools sort -n DJB05_th2out/accepted_hits.bam -o DJB05_sortname.bam
$ samtools view -o DJB05_sortname.sam DJB05_sortname.bam
#################################################################
# COUNT READS USING HTSEQ-COUNT
$ htseq-count -s no -a 10 DJB05_sortname.sam genes.gtf > DJB05.count
#################################################################
# Count-based differential analysis with edgeR
$ module load R
$ cd ∼/data/2016_rerun_myrnaseq/
$ Rscript edgeR_filter1cpm_updated.R GSad_2016.filelist
#################################################################
#Below is script within “edgeR_filter1cpm_updated.R”
#!/usr/bin/Rscript
#structure of file:1 col of batch names, 2 columns of sample names, with label at top of each column, tab separated
#label1label2
#b1 samp1 samp4
#b2 samp2 samp5
#b3 samp3 samp6
#samples in same row are assumed to be in same batch args <-commandArgs(TRUE)
filename=args[1]
bampath=“∼/data/2016_rerun_myrnaseq”
annotation=“∼/data/2016_rerun_myrnaseq/genes.gtf”
baseoutdir=“∼/data/2016_rerun_myrnaseq/”
library(edgeR)
x=read.table(filename,header=T)
label1=colnames(x[1])
label2=colnames(x[2])
samplelist=c(as.vector(x[,1]),as.vector(x[,2]))
conditions=c(rep(label1,nrow(x)),rep(label2,nrow(x)))
batch=rep(row.names(x),2)
names(conditions)=samplelist
#convert condition and batch to factors–prob not necessary for DESeq but edgeR likes it condition=factor(conditions)
batch=factor(batch)
#set up output directory for this experiment, create if it doesn’t exist outdir=sprintf(“%s/%s-%s”,baseoutdir,label1,label2)
dir.create(outdir)
#read in count table
count.table=read.table(sprintf(“%s/%s-%s-rawcounts.txt”,outdir,label1,label2),header=T)
# filtering–keep only reads with > 1cpm in at least half the samples keep_cpm <-rowSums(cpm(count.table)>1) >=nrow(x)
keep_quantile <-rowSums(count.table)>quantile(rowSums(count.table), probs=.5)
#save output of cpm vs quantile filters to log file sink(sprintf(“%s/edgeR.log”,outdir),append=T,split=T)
cat(“Comparison table of cpm vs quantile filters. CPM>2 in half of samples, quantile at 50%.\n”) addmargins(table(keep_cpm, keep_quantile))
sink()
count.table <-count.table[keep_cpm, ]
edesign=model.matrix(∼batch+condition)
e <-DGEList(counts=count.table)
e <- calcNormFactors(e)
e <- estimateGLMCommonDisp(e, edesign)
e <-estimateGLMTrendedDisp(e, edesign)
e <- estimateGLMTagwiseDisp(e, edesign)
#print size factors to log file sink(sprintf(“%s/edgeR.log”,outdir),append=T,split=T) cat(“Normalization factors:\n”)
e$samples sink()
#print dispersion and PCA plots to pdf
pdf(sprintf(“%s/%s-%s-edgeRdispersion.pdf”,outdir,label1,label2))
plotBCV(e, cex=0.4, main=“edgeR: Biological coefficient of variation (BCV) vs abundance”) dev.off()
pdf(sprintf(“%s/%s-%s-edgeRPCA.pdf”,outdir,label1,label2)) plotMDS(e, main=“edgeR MDS Plot”)
dev.off()
#Fit curves to GLM
efit <- glmFit(e, edesign)
efit <- glmLRT(efit, coef=sprintf(“condition%s”,label1)) #make results table and save
stats <- topTags(efit, n=nrow(e))$table
cpms <- cpm(e,)[rownames(stats),normalized.lib.sizes=T] etable=data.frame(stats,cpms)
etable <- etable[order(etable$FDR), ]
write.table(etable,file=sprintf(“%s/%s-%s-edgeR-filtered.txt”,outdir,label1,label2),quote=F,sep=“\t”) #copy sample file (argument) to outdir
file.copy(filename,sprintf(“%s/%s-%s-samplelist”,outdir,label1,label2))
#################################################################
#Below is script within “GSad_2016.filelist”
GSadPos GSadNeg b1 DJB07 DJB08
b2 DJB30 DJB31
b3 DJB28 DJB29
b4 DJB16 DJB21
b5 DJB36 DJB37
b6 DJB38 DJB39
#################################################################
~~~

#### Antibodies for immunohistochemistry

For these studies, the following primary antibodies were used: rabbit anti-melanopsin (Advanced Targeting Systems; 1:10,000), guinea pig anti-RBPMS (PhosphoSolutions 1832-RBPMS), rabbit anti-green fluorescent protein (GFP; Invitrogen); Goat anti-Brn3b antibody (Santa Cruz #sc-6026); mouse anti-Rasgrp1 (Santa Cruz sc-8430); guinea pig anti-Tbx20[1:8500] (Song et al., 2006)Rabbit anti-Giantin (Abcam ab24586). Secondary antibodies consisted of Alexa Fluor 350, 488, 594 or 647 donkey anti-goat, Alexa Fluor 594 donkey anti-rabbit and Alexa Fluor 594 goat anti-guinea pig.

#### Retina Tissue Preparations and Solutions

Mice were euthanized by inhalation of CO2. Prior to removing the eye, the dorsal margin of the cornea was marked with a cautery and this was used to guide the placement of a large relieving cut in the dorsal retina as a subsequent guide to retinal orientation. Eyes were removed immediately after death and placed in Hibernate-A solution preheated to 37 °C. To keep track of retinal orientation, the right and left eye were identified and processed separately.

#### Immunohistochemistry

After the retina was removed from the eye, it was placed on Millipore nitrocellulose paper. The retinas were fixed for 30 minutes at room temperature using 4% paraformaldehyde freshly prepared in 0.1M phosphate buffered saline (PBS; pH 7.4). The tissue was then washed for 15 minutes in PBS three times. The tissue was then incubated in a blocking solution of 0.5% Triton-X and 5% Goat Serum in PBS for two hours at room temperature. The tissue was incubated for two nights at 4 °C while on a shaker in the primary antibodies diluted in this same blocking solution. The following day, the samples were washed six times for 20 minutes in 0.1% Tween-20 in PBS. The tissue was then incubated for two hours in the appropriate Invitrogen or Jackson labs secondary antibodies diluted 1:1000 in the blocking solution at room temperature. The tissue was then washed six times for 10 minutes in 0.1% Tween-20 in PBS. The retinas were then mounted in Aquamount, coverslipped, and sealed with fingernail polish.

For Rasgrp1 immunofluorescence studies, an additional antigen retrieval step was included. antigen retrieval the tissue was then placed in Tris-EDTA (pH 8.0) for 30 minutes at 80 °C. The samples were then allowed to return to room temperatures (about 15-30 minutes) before they were removed from the Tris-EDTA solution and washed three times for 15 minutes in PBS.

#### Image Acquisition

Immunofluorescent images were captured on a Zeiss Confocal (LSM 510) and Nikon Eclipse microscope (Micro Video Instruments, Inc. E614, Avon, MA) with a built in Spot Camera (Diagnostic Instruments, Inc. HRD 100-NIK Sterling Heights, MI). Confocal images were taken with a 20x objective (Plan Apochromat, WD 0.55 mm) at a resolution of 2048 pixels. To enhance clarity, image files were pseudocolored and the brightness and contrast was adjusted using ImageJ 1.47 (National Insistute of Heath, Bethesda, MD). All final images were constructed using ImageJ and Powerpoint (Microsoft Corporation, Redmond, WA).

#### Analysis of Rasgrp1 expression in ipRGC subtypes

Because M1 and M2 cells have the highest levels of melanopsin expression of the ipRGC population, their dendrites were clearly visible and decipherable with immunostaining. M1 cells were identified by their dendritic projections to the OFF layer of the inner plexiform layer (IPL). M2 cells were identified by their dendrites which monostratify the ON sublamina of the IPL (Berson et al. 2010; Schmidt and Kofuji 2009). M3 cells bistratify the ON and OFF sublamina of the IPL (Schmidt and Kofuji 2011) and because these cells have similar levels of melanopsin expression and soma size as M2 cells, it is likely that M3 cells were included in the M2 population quantified in this study.

M4-M6 cells have the lowest levels of melanopsin-expression of the ipRGC population and their dendrites were not visible or decipherable with immunostaining. However, at least some members of the M4/M5/M6 population had lightly immunoreactive somas (Ecker et al., 2010; Quattrochi et al., 2018; Stabio et al., 2018). M4 cells were identified by their large soma size, low melanopsin immunodetectability and the lack of dendritic labeling (Estevez et al., 2012). The M5 and M6 cells we observed were identified by their low levels of melanopsin labeling and M2-sized somas (Ecker et al., 2010; Quattrochi et al., 2018; Stabio et al., 2018).

#### Analysis of Tbx20 expression in Cdh3-GFP mice

ipRGC subtypes were identified using a combination of morphological clues and process of elimination. In the case of M1 and M2 cells, which express the highest levels of melanopsin, confocal images of their immunofluorescence reveal dendritic information. As a result, unique dendritic features distinguish M1 and M2 cells. Cells with dendrites stratifying in the OFF layer of the IPL were identified as M1 cells, whereas cells stratifying in the ON layer were identified as M2 cells (Berson et al. 2010; Schmidt and Kofuji 2009). Using this method, M3 cells, which stratify in the ON and OFF sublamina of the IPL, were included in the M1 cell population unless otherwise noted (Schmidt and Kofuji 2011). As a result of their low levels of melanopsin, M4 and M5/6 cells are weakly labeled using anti-melanopsin immunohistochemistry with only some of their somata, but no dendrites, visible (Ecker et al., 2010; Estevez et al., 2012; Quattrochi et al., 2018; Stabio et al., 2018). Therefore, the method employed for deciphering M1 and M2 cells cannot be used to identify M4 and M5/6 cells. Instead, M4 cells were categorized by their lack of dendritic labeling, and their large soma size (Estevez et al., 2012). M5/6 cells were characterized by their lack of dendritic labeling, M2-sized somata, and process of elimination (Ecker et al., 2010; Estevez et al., 2012; Quattrochi et al., 2018; Stabio et al., 2018). In other words, cells that were not stained by the melanopsin antibody (which would detect M1-M3), but were labeled by GFP (labeling M1-M6), and had small somata, fell into the M5/6 category.

#### Brain injection of retrobeads into suprachiasmatic nucleus

Adult wild type mice (P30-P60) were anaesthetized with isofluorance and fluorescently labelled rhodamine latex microspheres (RetroBeads, Lumafluor) were injected into the ipsilateral suprachiasmatic nucleus to retrogradely label RGCs with axon terminals at the injections site. Three to five days later, the brain was removed and immediately fixed overnight. The following day, the brain was rinsed in 0.1M PBS and sectioned at 50um in the coronal plane. The slices were incubated with DAPI staining for 10 minutes was done in order to provide a reference for SCN location as indicated by concentrated cellular staining at the SCN. The slices were imaged for DAPI in UV channel and overlayed with rhodamine channel to identify the injection site in relation to SCN location. Special attention was paid to ensure that the injection did not extend into optic nerve and confound results by introducing off-target RGC labeling of fibers of passage.

#### Brain histology

Animals were sacrificed via transcardial perfusion, and brains were removed and incubated in 4% paraformaldehyde overnight. Brains were sectioned at 50μm in the coronal plane. To reveal individual processes in viral tracing experiments, virallyexpressed EYFP was enhanced using rabbit-anti-gfp (1:1000) and goat-anti-rabbit alexa 488 (1:500). To observe parvalbumin expression in OPN neurons, parvalbumin immunohistochemistry was performed using mouse-anti-parvalbumin (1:1000) followed by goat-anti-mouse alexa 488.

Stained and sliced brain slices were mounted on glass cover slides and imaged using a SPOT RT Slider digital microscope camera mounted to a Nikon (Diagnostic Instruments) as described previously (Berson et al., 2010; Estevez et al., 2012). Images were assembled in Adobe Photoshop CS3.

---

**Figure 1—figure supplement 1.**
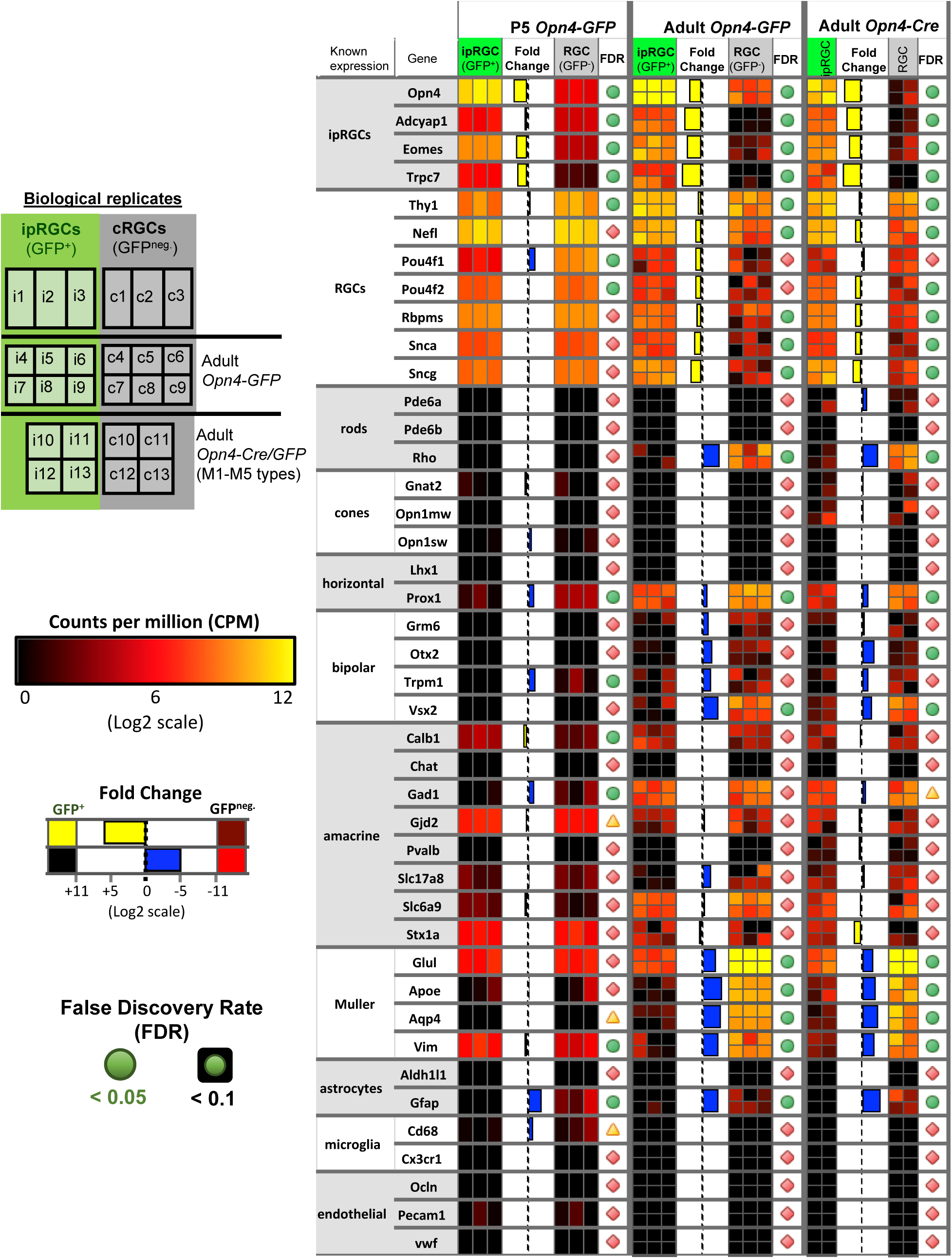
Purity and cell composition assessment of ipRGC and generic RGC samples. Heat map of known cell type marker gene expression in the retina to assess purity and cell composition of ipRGC and generic RGC samples. Shown are biological replicates tested for *Opn4-GFP* (P5 and adult) and *Opn4-Cre/GFP* reporters. Relative expression levels, fold change, and false discovery rate (FDR) are color-coded as indicated in the figure. White boxes indicate high gene expression, while blue represents little or no detected expression. FDR is not available (“NA”) in cases that our analysis filtered out genes with very low counts, less than 1 count per million (cpm), in more than half of the samples used in the differential expression analysis.

**Figure 2—figure supplement 1.**
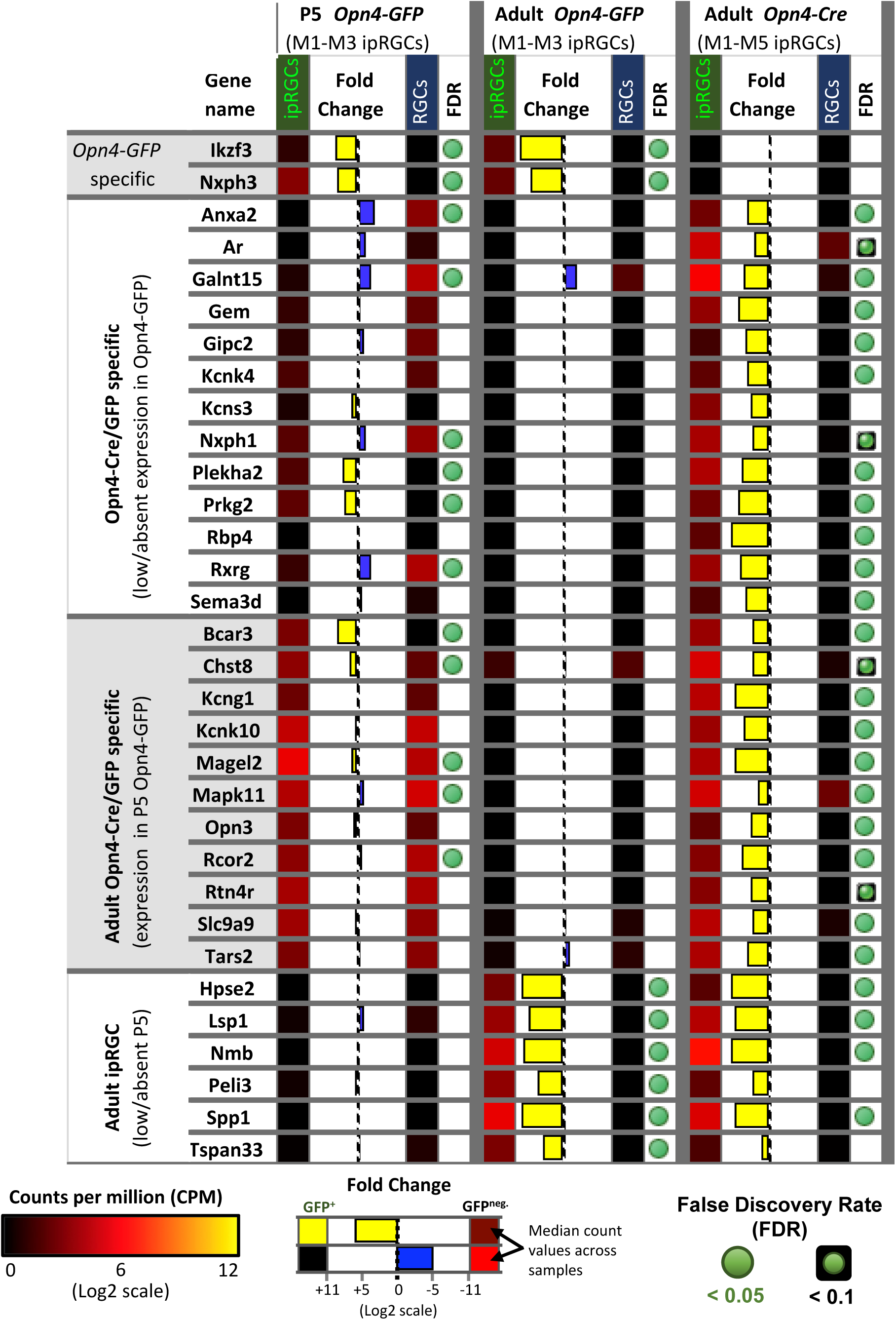
Heat map of genes differentially expressed in adult ipRGCs labeled by the *Opn4-Cre/GFP* reporter (M1-M6 ipRGCs) compared to *Opn4-GFP* (M1-M3 ipRGCs). Relative expression levels, fold change, and FDR are color-coded as indicated in the figure.

**Figure 2—figure supplement 2.**
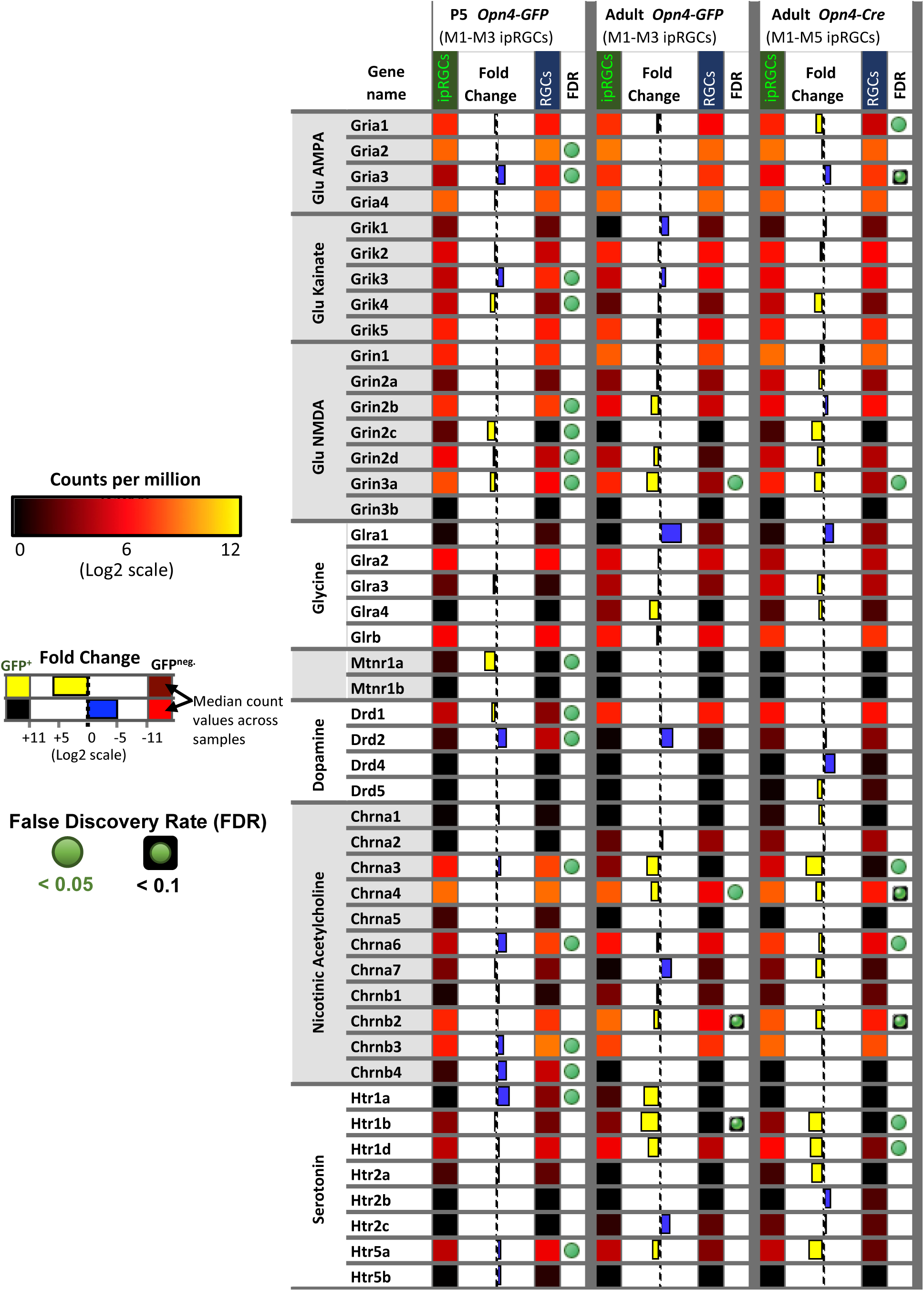
Heat map of genes encoding for nicotinic acetylcholine, dopamine, serotonin, glycine, glutamate, and melatonin receptors. Relative expression levels, fold change, and FDR are color-coded as indicated in the figure.

**Figure 2—figure supplement 3.**
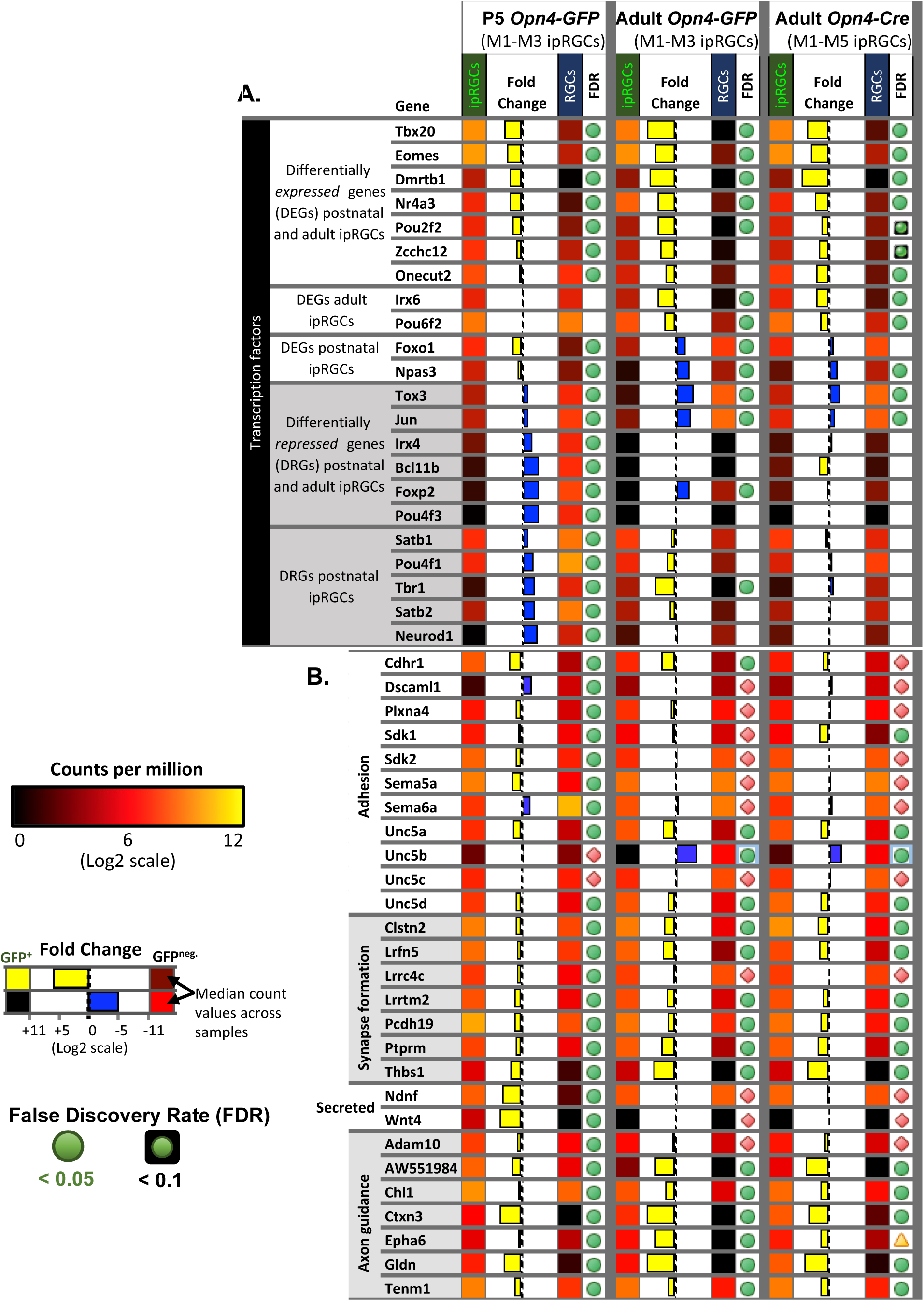
The expression pattern of developmentally regulated genes in ipRGCs. A. Heat map of genes encoding transcription factors that have a particular temporal pattern of differential expression in ipRGCs (e.g., high gene expression in P5 ipRGCs relative to adult expression). Relative expression levels, fold change, and FDR are color-coded as indicated in the figure. B. Heat map of genes relevant for development of ipRGCs. Relative expression levels, fold change, and FDR are color-coded as indicated in the figure.

**Figure 2—figure supplement 4.**
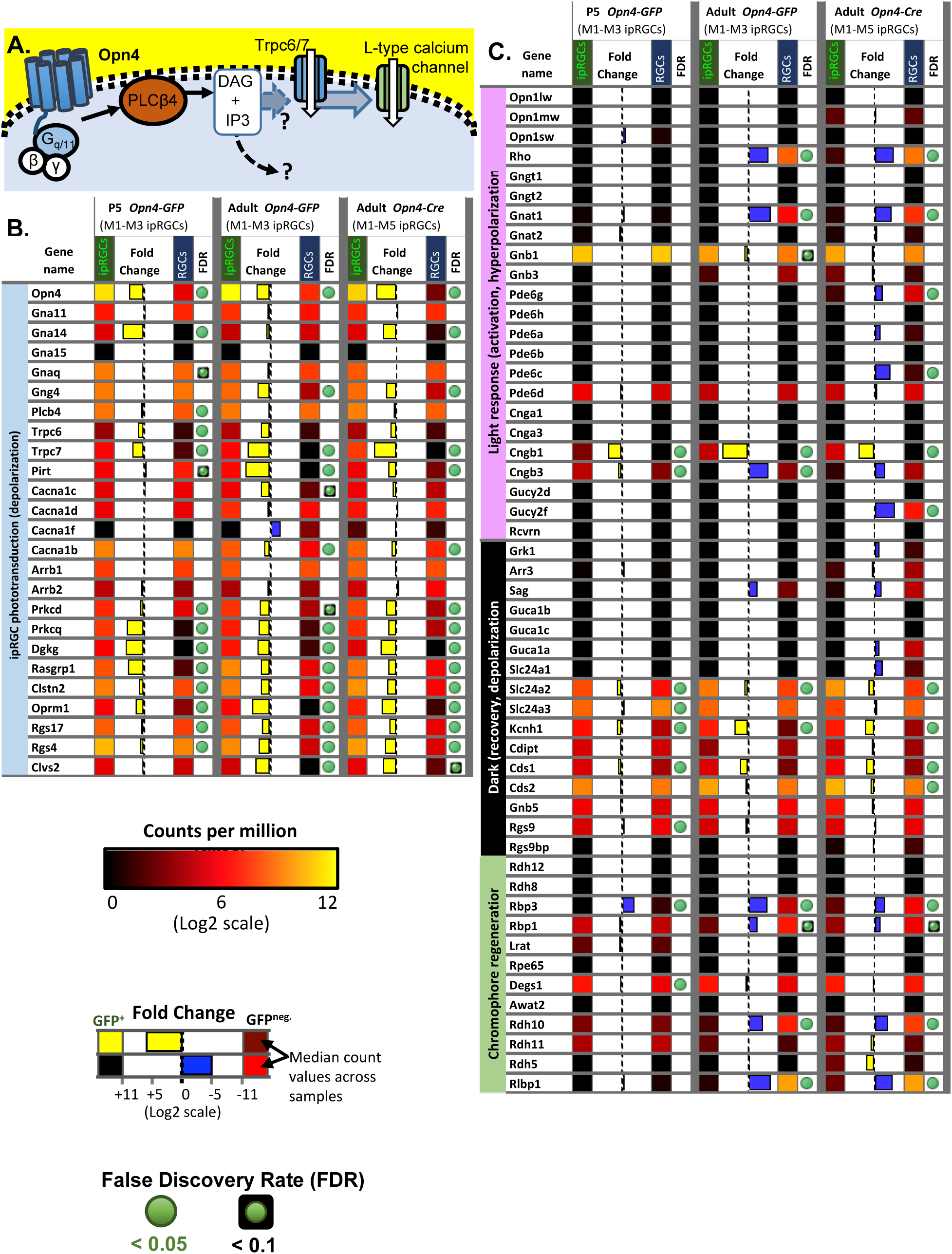
A. Distinct from rod and cone photoreceptors, the light-activation of Opn4 triggers a membrane-bound signaling cascade including G_q/11_ type G-proteins, the generation of 1,2-diacylglycerol (DAG) by PLCβ4, the opening of downstream TRPC6 and TRPC7 channels, and ultimately leads to the influx of calcium through L-type voltage-gated calcium channels. B. Heat map of genes that are potentially relevant to the Opn4-mediated phototransduction signaling cascade. Relative expression levels, fold change, and FDR are color-coded as indicated in the figure. C.Heat map of genes previously described to play a role in the light response, dark adaptation, and chromophore regeneration of rod and cone photoreceptors. Relative expression levels, fold change, and FDR are color-coded as indicated in the figure.

**Figure 3—figure supplement 1.**
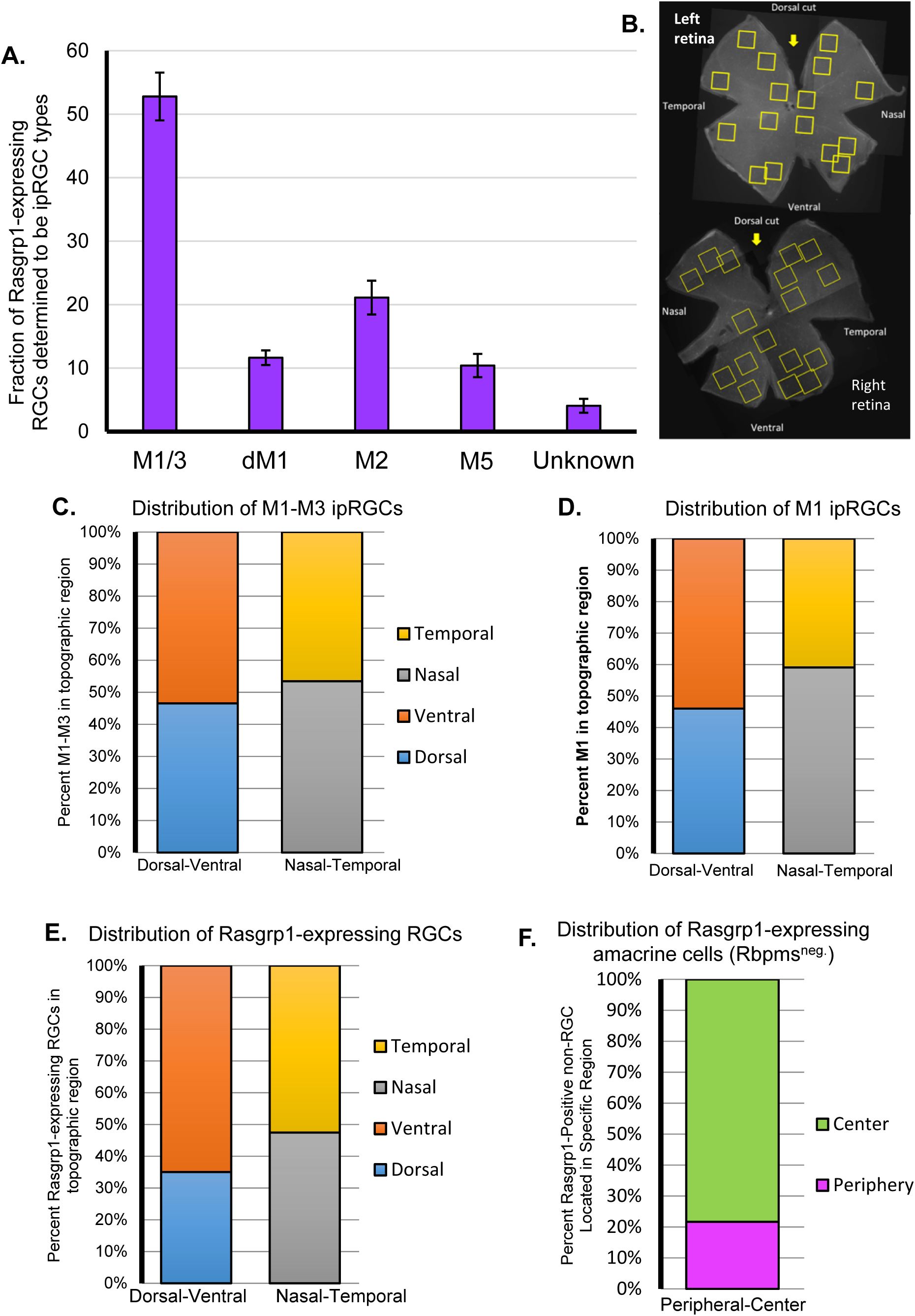
A.Distribution of all Rasgrp1-expressing RGCs (Rasgrp1^+^;Rbpms^+^) that belong to specific RGC types, to the extent that could be determined, including Opn4-immunoreactive ipRGC subtypes. No examples of M4 cells were observed to express Rasgrp1. The vast majority (96%) of Rasgrp1-RGCs are Opn4-immunopositive and therefore ipRGCs. The remaining “unknown” RGC types expressing Rasgrp1 (Rasgrp1^+^; Rbpms^+^; Opn4^neg.^) could be a low-expressing ipRGC type or conventional RGCs. Error bars represent standard error of the mean. B. Areas sampled for two wholemount wild type retinas immunostained for Rasgrp1, Opn4 and RBPMS. Yellow squares represent the locations of confocal images used for cell quantification. C. Analysis of the M1-M3 ipRGC population did not suggest a gradient of ipRGC spatial distribution across the retina. Seven frames were used to represent each area of the retina to maintain equal spatial contribution. Total M1-M3 represented in the dorsal-ventral and nasal-temporal columns is 217. E. Analysis of the M1 cell population revealed a slight naso-temporal gradient of M1 spatial distribution across the retina. Seven frames were used to represent each area of the retina to maintain equal spatial contribution. Total M1s represented in the dorsal-ventral column and nasal-temporal columns is 113 and 110, respectively. F. Analysis of the Rasgrp1-positive RGC population suggests a slight ventral-dorsal gradient of Rasgrp1-positive RGCs spatial distribution across the retina. Seven frames were used to represent each area of the retina to maintain equal spatial contribution. Total Rasgrp1-positive RGCs represented in the dorsal-ventral column and nasal-temporal column is 117 and 118, respectively. F. The amacrine cells (presumed) expressing Rasgrp1 exhibited a dramatic center-peripheral gradient of Rasgrp1-positive non-RGC spatial distribution across the retina. Four frames were used to represent each area of the retina to maintain equal spatial contribution. The total Rasgrp1-positive non-RGCs represented in the periphery and center was 314.

**Figure 4—figure supplement 1.**
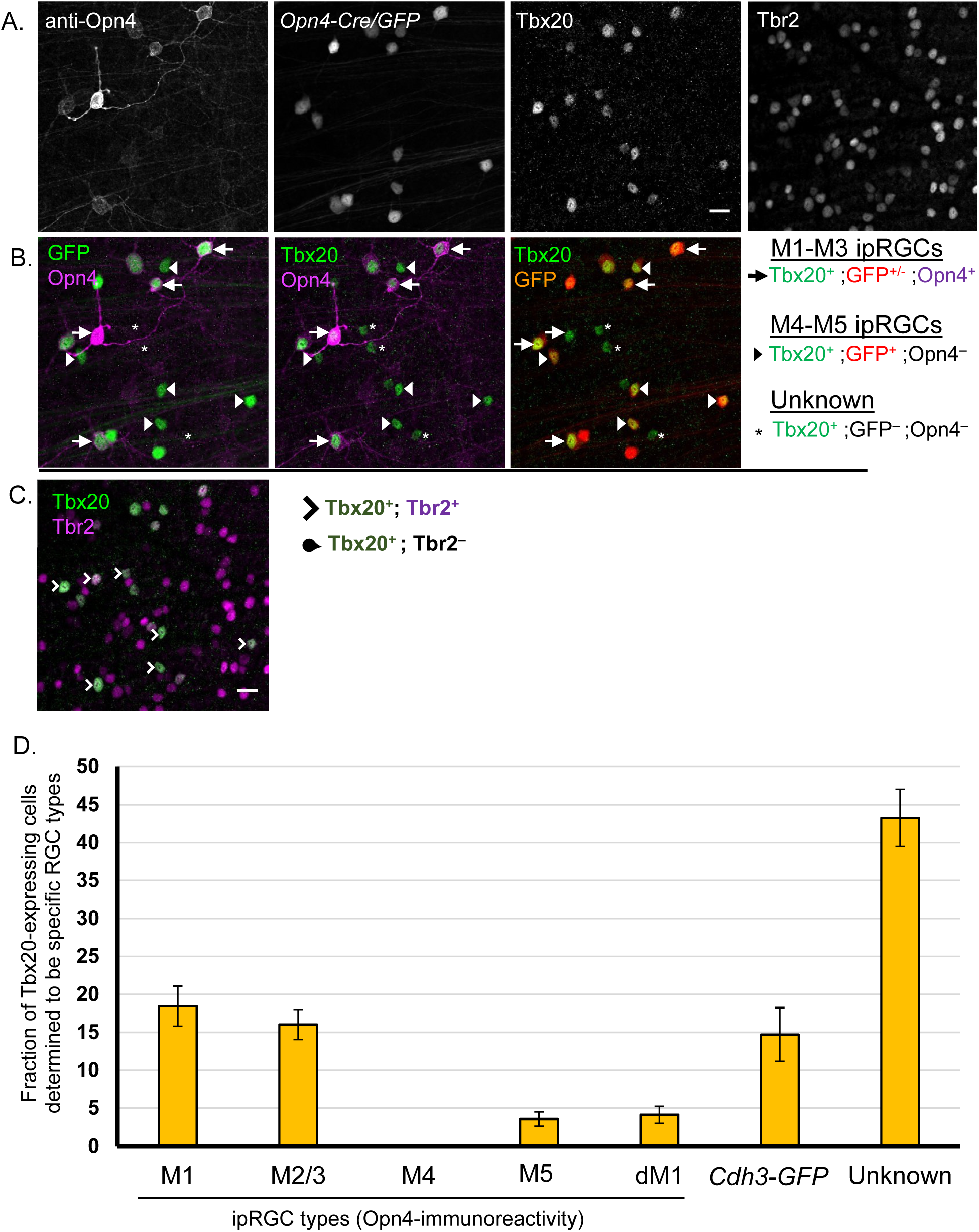
Coexpression study of Tbx20 with Opn4-Cre/GFP and Tbr2, including distribution of Tbx20-expression across ipRGC subtypes. A-C. Quadruple immunofluorescence of Opn4, Tbx20, Opn4-Cre/GFP, and Tbr2. Scale bar, 20 μm. A. Gray scale of Opn4, Opn4-Cre/GFP, and Tbx20 immunofluorescence. B. Co-expression study of Tbx20 (green) in the context of Opn4 (magenta) and *Opn4-Cre/GFP* (red) labeling. GFP cells that are Opn4-immunonegative are inferred M4-M6 types. C. Co-expression analysis of Tbr2 (magenta) with Tbx20 (green). D. Distribution of Tbx20 expressing cells that belong to specific RGC types, to the extent that could be determined, including Opn4-immunoreactive ipRGC subtypes and RGCs labeled by the *Cdh3-GFP* transgenic reporter. Unaccounted Tbx20-expressing cells are designated as “unknown” RGC types. Error bars represent standard error of the mean.

**Figure 5—figure supplement 1.**
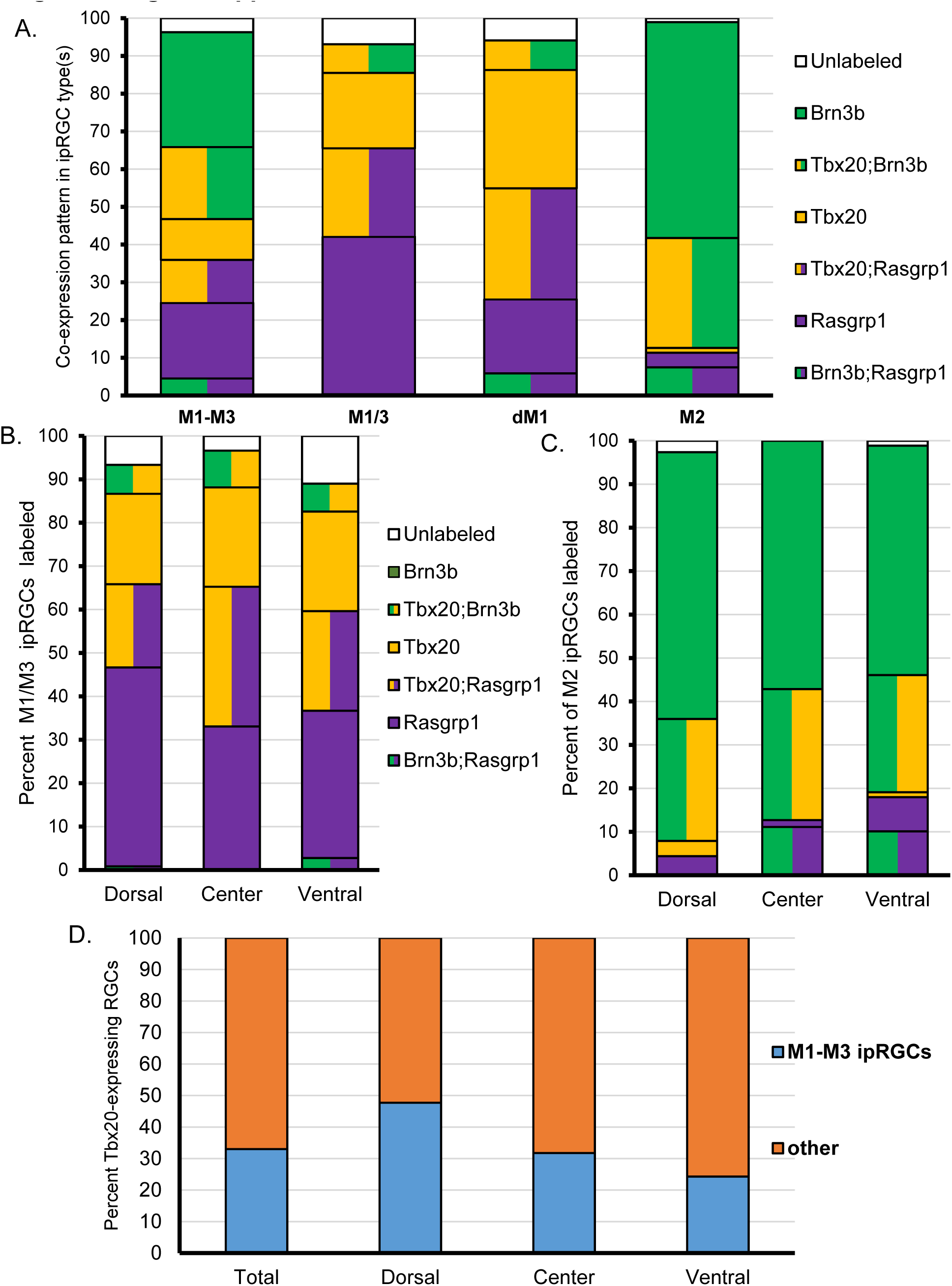
Topographic distribution and ipRGC subtype-specific quantification of Rasgrp1-Tbx20-Brn3b expression. A.Comparison of co-expression patterns of Brn3b, Rasgrp1, and Tbx20 within group of combined M1-M3 (M1+M2+M3) ipRGCs, the M1/3 ipRGCs (M1+M3), displaced M1s, and M2 ipRGCs. B. Lack of major topographic variations in the fraction of M1/M3 ipRGCs immunoreactive for Rasgrp1, Tbx20, and Brn3b. C. Comparison of Rasgrp1-Tbx20-Brn3b expression pattern in M2 ipRGCs across dorsal, center, and ventral regions of the retina. D. Distribution of Tbx20-expression in M1-M3 ipRGCs compared to Opn4-immunonegative cells in the context of topgraphic regions across the retina.

## References

Anders S, Huber W (2010) Differential expression analysis for sequence count data. Genome Biology 2010 11:10 11:R106.

Anders S, McCarthy DJ, Chen Y, Okoniewski M, Smyth GK, Huber W, Robinson MD (2013) Count-based differential expression analysis of RNA sequencing data using R and Bioconductor. Nature Protocols 8:1765–1786.

Atlasz T, Szabadfi K, Kiss P, Racz B, Gallyas F, Tamas A, Gaal V, Marton Z, Gabriel R, Reglodi D (2010) Pituitary adenylate cyclase activating polypeptide in the retina: focus on the retinoprotective effects. Ann N Y Acad Sci 1200:128–139.

Barres BA, Silverstein BE, Corey DP, Chun LLY (1988) Immunological, morphological, and electrophysiological variation among retinal ganglion cells purified by panning. Neuron 1:791–803.

Bedont JL, Blackshaw S (2015) Constructing the suprachiasmatic nucleus: a watchmaker’s perspective on the central clockworks. Front Syst Neurosci 9:74.

Beglopoulos V, Montag-Sallaz M, Rohlmann A, Piechotta K, Ahmad M, Montag D, Missler M (2005) Neurexophilin 3 is highly localized in cortical and cerebellar regions and is functionally important for sensorimotor gating and motor coordination. Mol Cell Biol 25:7278–7288.

Berson DM, Castrucci AM, Provencio I (2010) Morphology and mosaics of melanopsinexpressing retinal ganglion cell types in mice. Journal of Comparative Neurology 518:2405–2422.

Bivona TG, Pérez De Castro I, Ahearn IM, Grana TM, Chiu VK, Lockyer PJ, Cullen PJ, Pellicer A, Cox AD, Philips MR (2003) Phospholipase Cgamma activates Ras on the Golgi apparatus by means of RasGRP1. Nature 424:694–698.

Brewer GJ (1997) Isolation and culture of adult rat hippocampal neurons. Journal of Neuroscience Methods 71:143–155.

Brewer GJ, Torricelli JR (2007) Isolation and culture of adult neurons and neurospheres. Nature Protocols 2:1490–1498.

Brewer GJ, Torricelli JR, Evege EK, Price PJ (1993) Optimized survival of hippocampal neurons in B27-supplemented neurobasal(tm), a new serum-free medium combination. Journal of Neuroscience Research 35:567–576.

Burden-Gulley SM, Brady-Kalnay SM (1999) PTPmu regulates N-cadherin-dependent neurite outgrowth. J Cell Biol 144:1323–1336.

Cahoy JD, Emery B, Kaushal A, Foo LC, Zamanian JL, Christopherson KS, Xing Y, Lubischer JL, Krieg PA, Krupenko SA, Thompson WJ, Barres BA (2008) A transcriptome database for astrocytes, neurons, and oligodendrocytes: a new resource for understanding brain development and function. J Neurosci 28:264–278.

Cai C-L, Zhou W, Yang L, Bu L, Qyang Y, Zhang X, Li X, Rosenfeld MG, Chen J, Evans S (2005) T-box genes coordinate regional rates of proliferation and regional specification during cardiogenesis. Development 132:2475–2487.

Cameron EG, Robinson PR (2014) β-Arrestin-Dependent Deactivation of Mouse Melanopsin Craft CM, ed. PLoS ONE 9:e113138.

Caretti E, Devarajan K, Coudry R, Ross E, Clapper ML, Cooper HS, Bellacosa A (2008) Comparison of RNA amplification methods and chip platforms for microarray analysis of samples processed by laser capture microdissection. Journal of Cellular Biochemistry 103:556–563.

Carson CT, Kinzler ER, Parr BA (2000) Tbx12, a novel T-box gene, is expressed during early stages of heart and retinal development. Mechanisms of Development 96:137–140.

Carson CT, Pagratis M, Parr BA (2004) Tbx12 regulates eye development in Xenopus embryos. Biochemical and Biophysical Research Communications 318:485–489.

Chen SK, Badea TC, Hattar S (2011) Photoentrainment and pupillary light reflex are mediated by distinct populations of ipRGCs. Nature 476:92–95.

Clément-Ziza M, Gentien D, Lyonnet S, Thiery J-P, Besmond C, Decraene C (2009) Evaluation of methods for amplification of picogram amounts of total RNA for whole genome expression profiling. BMC Genomics 10:246.

Craig AM, Kang Y (2007) Neurexin-neuroligin signaling in synapse development. Current Opinion in Neurobiology 17:43–52.

Cui Q, Ren C, Sollars PJ, Pickard GE, So KF (2015) The injury resistant ability of melanopsin-expressing intrinsically photosensitive retinal ganglion cells. Neuroscience 284:845–853.

de Sevilla Müller LP, Sargoy A, Rodriguez AR, Brecha NC (2014) Melanopsin Ganglion Cells Are the Most Resistant Retinal Ganglion Cell Type to Axonal Injury in the Rat Retina Tosini G, ed. PLoS ONE 9:e93274.

de Wit J, Sylwestrak E, O’sullivan ML, Otto S, Tiglio K, Savas JN, Yates JR, Comoletti D, Taylor P, Ghosh A (2009) LRRTM2 interacts with Neurexin1 and regulates excitatory synapse formation. Neuron 64:799–806.

Diaz A, Ruiz F, Flórez J, Hurlé MA, Pazos A (1995) Mu-opioid receptor regulation during opioid tolerance and supersensitivity in rat central nervous system. J Pharmacol Exp Ther 274:1545–1551.

Doğrul A, Yeşilyurt O, Işimer A, Güzeldemir ME (2001) L-type and T-type calcium channel blockade potentiate the analgesic effects of morphine and selective mu opioid agonist, but not to selective delta and kappa agonist at the level of the spinal cord in mice. Pain 93:61–68.

Dower NA, Stang SL, Bottorff DA, Ebinu JO, Dickie P, Ostergaard HL, Stone JC (2000) RasGRP is essential for mouse thymocyte differentiation and TCR signaling. Nat Immunol 1:317–321.

Duan X, Qiao M, Bei F, Kim I-J, He Z, Sanes JR (2015) Subtype-Specific Regeneration of Retinal Ganglion Cells following Axotomy: Effects of Osteopontin and mTOR Signaling. Neuron 85:1244–1256.

Dumitrescu ON, Pucci FG, Wong KY, Berson DM (2009) Ectopic retinal ON bipolar cell synapses in the OFF inner plexiform layer: Contacts with dopaminergic amacrine cells and melanopsin ganglion cells. Journal of Comparative Neurology 517:226–244.

Ebinu JO, Bottorff DA, Chan EY, Stang SL, Dunn RJ, Stone JC (1998) RasGRP, a Ras guanyl nucleotide-releasing protein with calcium- and diacylglycerol-binding motifs. Science 280:1082–1086.

Ecker JL, Dumitrescu ON, Wong KY, Alam NM, Chen S-K, LeGates T, Renna JM, Prusky GT, Berson DM, Hattar S (2010) Melanopsin-Expressing Retinal Ganglion-Cell Photoreceptors: Cellular Diversity and Role in Pattern Vision. Neuron 67:49–60.

Eide EJ, Woolf MF, Kang H, Woolf P, Hurst W, Camacho F, Vielhaber EL, Giovanni A, Virshup DM (2005) Control of mammalian circadian rhythm by CKIepsilon-regulated proteasome-mediated PER2 degradation. Mol Cell Biol 25:2795–2807.

Emanuel AJ, Do MTH (2015) Melanopsin Tristability for Sustained and Broadband Phototransduction. Neuron 85:1043–1055.

Emanuel AJ, Kapur K, Do MTH (2017) Biophysical Variation within the M1 Type of Ganglion Cell Photoreceptor. Cell Rep 21:1048–1062.

Estevez ME, Fogerson PM, Ilardi MC, Borghuis BG, Chan E, Weng S, Auferkorte ON, Demb JB, Berson DM (2012) Form and function of the M4 cell, an intrinsically photosensitive retinal ganglion cell type contributing to geniculocortical vision. J Neurosci 32:13608–13620.

Fernandez DC, Chang Y-T, Hattar S, Chen S-K (2016) Architecture of retinal projections to the central circadian pacemaker. Proc Natl Acad Sci USA 113:6047–6052.

Fink M, Lesage F, Duprat F, Heurteaux C, Reyes R, Fosset M, Lazdunski M (1998) A neuronal two P domain K+ channel stimulated by arachidonic acid and polyunsaturated fatty acids. EMBO J 17:3297–3308.

Fornaro M, Raimondo S, Lee JM, Giuseppina Giacobini-Robecchi M (2007) Neuron-specific Hu proteins sub-cellular localization in primary sensory neurons. Annals of Anatomy - Anatomischer Anzeiger 189:223–228.

Frings S, Brüll N, Dzeja C, Angele A, Hagen V, Kaupp UB, Baumann A (1998) Characterization of ether-à-go-go channels present in photoreceptors reveals similarity to IKx, a K+ current in rod inner segments. J Gen Physiol 111:583–599.

Gorentla BK, Wan C-K, Zhong X-P (2011) Negative regulation of mTOR activation by diacylglycerol kinases. Blood 117:4022–4031.

Graham DM, Wong KY, Shapiro P, Frederick C, Pattabiraman K, Berson DM (2008) Melanopsin Ganglion Cells Use a Membrane-Associated Rhabdomeric Phototransduction Cascade. Journal of Neurophysiology 99:2522–2532.

Haeryfar SMM, Hoskin DW (2004) Thy-1: More than a Mouse Pan-T Cell Marker. The Journal of Immunology 173:3581–3588.

Hannibal J, Hindersson P, Østergaard J, Georg B, Heegaard S, Larsen PJ, Fahrenkrug J (2004) Melanopsin Is Expressed in PACAP-Containing Retinal Ganglion Cells of the Human Retinohypothalamic Tract. Invest Ophthalmol Vis Sci 45:4202–4209.

Hartwick ATE, Bramley JR, Yu J, Stevens KT, Allen CN, Baldridge WH, Sollars PJ, Pickard GE (2007) Light-evoked calcium responses of isolated melanopsin-expressing retinal ganglion cells. J Neurosci 27:13468–13480.

Hattar S, Kumar M, Park A, Tong P, Tung J, Yau K-W, Berson DM (2006) Central projections of melanopsin-expressing retinal ganglion cells in the mouse. J Comp Neurol 497:326–349.

Heiman M, Schaefer A, Gong S, Peterson JD, Day M, Ramsey KE, Suárez-Fariñas M, Schwarz C, Stephan DA, Surmeier DJ, Greengard P, Heintz N (2008) A Translational Profiling Approach for the Molecular Characterization of CNS Cell Types. Cell 135:738–748.

Hinman MN, Lou H (2008) Diverse molecular functions of Hu proteins. Cell Mol Life Sci 65:3168–3181.

Hoshi H, Liu W-L, Massey SC, Mills SL (2009) ON inputs to the OFF layer: bipolar cells that break the stratification rules of the retina. J Neurosci 29:8875–8883.

Hu C, Hill DD, Wong KY (2013) Intrinsic physiological properties of the five types of mouse ganglion-cell photoreceptors. Journal of Neurophysiology 109:1876–1889.

Hughes S, Hankins MW, Foster RG, Peirson SN (2012) Melanopsin phototransduction: slowly emerging from the dark. Prog Brain Res 199:19–40.

Hughes S, Jagannath A, Hickey D, Gatti S, Wood M, Peirson SN, Foster RG, Hankins MW (2015) Using siRNA to define functional interactions between melanopsin and multiple G Protein partners. Cell Mol Life Sci 72:165–179.

Jain V, Ravindran E, Dhingra NK (2012) Differential expression of Brn3 transcription factors in intrinsically photosensitive retinal ganglion cells in mouse. Journal of Comparative Neurology 520:742–755.

Jeon C-J, Strettoi E, Masland RH (1998) The Major Cell Populations of the Mouse Retina. J Neurosci 18:8936–8946.

Ji M, Zhao W-J, Dong L-D, Miao Y, Yang X-L, Sun X-H, Wang Z (2011) RGS2 and RGS4 modulate melatonin-induced potentiation of glycine currents in rat retinal ganglion cells. Brain Research 1411:1–8.

Kim I-J, Zhang Y, Meister M, Sanes JR (2010) Laminar restriction of retinal ganglion cell dendrites and axons: subtype-specific developmental patterns revealed with transgenic markers. J Neurosci 30:1452–1462.

Koussounadis A, Langdon SP, Um IH, Harrison DJ, Smith VA (2015) Relationship between differentially expressed mRNA and mRNA-protein correlations in a xenograft model system. Sci Rep 5:10775.

Li S, Yang C, Zhang L, Gao X, Wang X, Liu W, Wang Y, Jiang S, Wong YH, Zhang Y, Liu K (2016) Promoting axon regeneration in the adult CNS by modulation of the melanopsin/GPCR signaling. Proc Natl Acad Sci USA 113:1937–1942.

Li S-Y, Yau S-Y, Chen B-Y, Tay DK, Lee VWH, Pu M-L, Chan HHL, So K-F (2008) Enhanced Survival of Melanopsin-expressing Retinal Ganglion Cells After Injury is Associated with the PI3 K/Akt Pathway. Cell Mol Neurobiol 28:1095–1107.

Lin JC, Ho W-H, Gurney A, Rosenthal A (2003) The netrin-G1 ligand NGL-1 promotes the outgrowth of thalamocortical axons. Nature Neuroscience 6:1270–1276.

Lin Z, Liu J, Ding H, Xu F, Liu H (2018) Structural basis of SALM5-induced PTPδ dimerization for synaptic differentiation. Nature Communications 9:268.

Lipina TV, Prasad T, Yokomaku D, Luo L, Connor SA, Kawabe H, Wang YT, Brose N, Roder JC, Craig AM (2016) Cognitive Deficits in Calsyntenin-2-deficient Mice Associated with Reduced GABAergic Transmission. Neuropsychopharmacology 41:802–810.

Lobo MK, Karsten SL, Gray M, Geschwind DH, Yang XW (2006) FACS-array profiling of striatal projection neuron subtypes in juvenile and adult mouse brains. Nature Neuroscience 9:443–452.

Macosko EZ, Basu A, Satija R, Nemesh J, Shekhar K, Goldman M, Tirosh I, Bialas AR, Kamitaki N, Martersteck EM, Trombetta JJ, Weitz DA, Sanes JR, Shalek AK, Regev A, McCarroll SA (2015) Highly Parallel Genome-wide Expression Profiling of Individual Cells Using Nanoliter Droplets. Cell 161:1202–1214.

Mao C-A, Kiyama T, Pan P, Furuta Y, Hadjantonakis A-K, Klein WH (2008) Eomesodermin, a target gene of Pou4f2, is required for retinal ganglion cell and optic nerve development in the mouse. Development 135:271–280.

Mao C-A, Li H, Zhang Z, Kiyama T, Panda S, Hattar S, Ribelayga CP, Mills SL, Wang SW (2014) T-box transcription regulator Tbr2 is essential for the formation and maintenance of Opn4/melanopsin-expressing intrinsically photosensitive retinal ganglion cells. J Neurosci 34:13083–13095.

Mao H, Zhao Q, Daigle M, Ghahremani MH, Chidiac P, Albert PR (2004) RGS17/RGSZ2, a novel regulator of Gi/o, Gz, and Gq signaling. J Biol Chem 279:26314–26322.

Martin S, Lino de Oliveira C, Mello de Queiroz F, Pardo LA, Stühmer W, Del Bel E (2008) Eag1 potassium channel immunohistochemistry in the CNS of adult rat and selected regions of human brain. Neuroscience 155:833–844.

Matsuoka RL, Nguyen-Ba-Charvet KT, Parray A, Badea TC, Chédotal A, Kolodkin AL (2011) Transmembrane semaphorin signalling controls laminar stratification in the mammalian retina. Nature 470:259–263.

Matsushima D, Heavner W, Pevny LH (2011) Combinatorial regulation of optic cup progenitor cell fate by SOX2 and PAX6. Development 138:443–454.

Missler M, Hammer RE, Südhof TC (1998) Neurexophilin binding to alpha-neurexins. A single LNS domain functions as an independently folding ligand-binding unit. J Biol Chem 273:34716–34723.

Moises HC, Rusin KI, Macdonald RL (1994) Mu- and kappa-opioid receptors selectively reduce the same transient components of high-threshold calcium current in rat dorsal root ganglion sensory neurons. J Neurosci 14:5903–5916.

Morrow EM, Belliveau MJ, Cepko CL (1998) Two Phases of Rod Photoreceptor Differentiation during Rat Retinal Development. J Neurosci 18:3738–3748.

Morse AM, Carballo V, Baldwin DA, Taylor CG, McIntyre LM (2010) Comparison between NuGEN’s WT-Ovation Pico and one-direct amplification systems. J Biomol Tech 21:141–147.

Naiche LA, Harrelson Z, Kelly RG, Papaioannou VE (2005) T-box genes in vertebrate development. Annu Rev Genet 39:219–239.

Novak A, Guo C, Yang W, Nagy A, Lobe CG (2000) Z/EG, a double reporter mouse line that expresses enhanced green fluorescent protein upon Cre-mediated excision. Genesis.

Oancea E, Meyer T (1998) Protein kinase C as a molecular machine for decoding calcium and diacylglycerol signals. Cell 95:307–318.

Pack W, Hill DD, Wong KY (2015) Melatonin modulates M4-type ganglion-cell photoreceptors. Neuroscience 303:178–188.

Pederick DT, Homan CC, Jaehne EJ, Piltz SG, Haines BP, Baune BT, Jolly LA, Hughes JN, Gecz J, Thomas PQ (2016) Pcdh19 Loss-of-Function Increases Neuronal Migration In Vitro but is Dispensable for Brain Development in Mice. Sci Rep 6:26765.

Peirson SN, Oster H, Jones SL, Leitges M, Hankins MW, Foster RG (2007) Microarray Analysis and Functional Genomics Identify Novel Components of Melanopsin Signaling. Current Biology 17:1363–1372.

Pettem KL, Yokomaku D, Takahashi H, Ge Y, Craig AM (2013) Interaction between autism-linked MDGAs and neuroligins suppresses inhibitory synapse development. J Cell Biol 200:321–336.

Pierret P, Dunn RJ, Djordjevic B, Stone JC, Richardson PM (2000) Distribution of ras guanyl releasing protein (RasGRP) mRNA in the adult rat central nervous system. J Neurocytol 29:485–497.

Pocock R, Mione M, Hussain S, Maxwell S, Pontecorvi M, Aslam S, Gerrelli D, Sowden JC, Woollard A (2008) Neuronal function of Tbx20 conserved from nematodes to vertebrates. Developmental Biology 317:671–685.

Puente LG, Stone JC, Ostergaard HL (2000) Evidence for protein kinase C-dependent and - independent activation of mitogen-activated protein kinase in T cells: potential role of additional diacylglycerol binding proteins. The Journal of Immunology 165:6865–6871.

Quail MA, Kozarewa I, Smith F, Scally A, Stephens PJ, Durbin R, Swerdlow H, Turner DJ (2008) A large genome center’s improvements to the Illumina sequencing system. Nat Methods 5:1005–1010.

Quattrochi LE, Stabio ME, Kim I, Berson DM (2018) The M6 cell: A small-field bistratified photosensitive ganglion cell. in press.

Reifler AN, Chervenak AP, Dolikian ME, Benenati BA, Li BY, Wachter RD, Lynch AM, Demertzis ZD, Meyers BS, Abufarha FS, Jaeckel ER, Flannery MP, Wong KY (2015) All Spiking, Sustained ON Displaced Amacrine Cells Receive Gap-Junction Input from Melanopsin Ganglion Cells. Current Biology 25:2763–2773.

Renna JM, Weng S, Berson DM (2011) Light acts through melanopsin to alter retinal waves and segregation of retinogeniculate afferents. Nature Neuroscience 14:827–829.

Rodriguez AR, Sevilla Müller LP, Brecha NC (2014) The RNA binding protein RBPMS is a selective marker of ganglion cells in the mammalian retina. Journal of Comparative Neurology 522:1411–1443.

Rousso DL, Qiao M, Kagan RD, Yamagata M, Palmiter RD, Sanes JR (2016) Two Pairs of ON and OFF Retinal Ganglion Cells Are Defined by Intersectional Patterns of Transcription Factor Expression. Cell Rep 15:1930–1944.

Sabbah S, Berg D, Papendorp C, Briggman KL, Berson DM (2017) A Cre Mouse Line for Probing Irradiance- and Direction-Encoding Retinal Networks. eNeuro 4:ENEURO.0065–17.2017.

Sakabe NJ, Aneas I, Shen T, Shokri L, Park S-Y, Bulyk ML, Evans SM, Nobrega MA (2012) Dual transcriptional activator and repressor roles of TBX20 regulate adult cardiac structure and function. Hum Mol Genet 21:2194–2204.

Sand A, Schmidt TM, Kofuji P (2012) Diverse types of ganglion cell photoreceptors in the mammalian retina. Progress in Retinal and Eye Research 31:287–302.

Sanes JR, Masland RH (2015) The Types of Retinal Ganglion Cells: Current Status and Implications for Neuronal Classification. http://dxdoiorg/101146/annurev-neuro-071714-034120 38:221–246.

Schmidt TM, Do MTH, Dacey D, Lucas R, Hattar S, Matynia A (2011) Melanopsin-positive intrinsically photosensitive retinal ganglion cells: from form to function. J Neurosci 31:16094–16101.

Schmidt TM, Taniguchi K, Kofuji P (2008) Intrinsic and Extrinsic Light Responses in Melanopsin-Expressing Ganglion Cells During Mouse Development. Journal of Neurophysiology 100:371–384.

Sengupta A, Baba K, Mazzoni F, Pozdeyev NV, Strettoi E, Iuvone PM, Tosini G (2011) Localization of melatonin receptor 1 in mouse retina and its role in the circadian regulation of the electroretinogram and dopamine levels. Yamazaki S, ed. PLoS ONE 6:e24483.

Shekhar K, Lapan SW, Whitney IE, Tran NM, Macosko EZ, Kowalczyk M, Adiconis X, Levin JZ, Nemesh J, Goldman M, McCarroll SA, Cepko CL, Regev A, Sanes JR (2016) Comprehensive Classification of Retinal Bipolar Neurons by Single-Cell Transcriptomics. Cell 166:1308–1323.e1330.

Shen T, Aneas I, Sakabe N, Dirschinger RJ, Wang G, Smemo S, Westlund JM, Cheng H, Dalton N, Gu Y, Boogerd CJ, Cai C-L, Peterson K, Chen J, Nobrega MA, Evans SM (2011) Tbx20 regulates a genetic program essential to adult mouse cardiomyocyte function. J Clin Invest 121:4640–4654.

Sheng W-L, Chen W-Y, Yang X-L, Zhong Y-M, Weng S-J (2015) Co-expression of two subtypes of melatonin receptor on rat M1-type intrinsically photosensitive retinal ganglion cells. Barnes S, ed. PLoS ONE 10:e0117967.

Shulga YV, Topham MK, Epand RM (2011) Regulation and Functions of Diacylglycerol Kinases. Chemical Reviews 111:6186–6208.

Siegert S, Cabuy E, Scherf BG, Kohler H, Panda S, Le Y-Z, Fehling HJ, Gaidatzis D, Stadler MB, Roska B (2012) Transcriptional code and disease map for adult retinal cell types. Nature Neuroscience 15:487–495.

Soto I, Oglesby E, Buckingham BP, Son JL, Roberson EDO, Steele MR, Inman DM, Vetter ML, Horner PJ, Marsh-Armstrong N (2008) Retinal Ganglion Cells Downregulate Gene Expression and Lose Their Axons within the Optic Nerve Head in a Mouse Glaucoma Model. J Neurosci 28:548–561.

Stabio ME, Sabbah S, Quattrochi LE, Ilardi MC, Fogerson PM, Leyrer ML, Kim MT, Kim I, Schiel M, Renna JM, Briggman KL, Berson DM (2018) The M5 Cell: A Color-Opponent Intrinsically Photosensitive Retinal Ganglion Cell. Neuron 97:251.

Star EN, Zhu M, Shi Z, Liu H, Pashmforoush M, Sauve Y, Bruneau BG, Chow RL (2012) Regulation of retinal interneuron subtype identity by the Iroquois homeobox gene Irx6. Development 139:4644–4655.

Stennard FA, Costa MW, Elliott DA, Rankin S, Haast SJP, Lai D, McDonald LPA, Niederreither K, Dolle P, Bruneau BG, Zorn AM, Harvey RP (2003) Cardiac T-box factor Tbx20 directly interacts with Nkx2-5, GATA4, and GATA5 in regulation of gene expression in the developing heart. Developmental Biology 262:206–224.

Sweeney NT, Tierney H, Feldheim DA (2014) Tbr2 is required to generate a neural circuit mediating the pupillary light reflex. J Neurosci 34:5447–5453.

Takeuchi JK, Mileikovskaia M, Koshiba-Takeuchi K, Heidt AB, Mori AD, Arruda EP, Gertsenstein M, Georges R, Davidson L, Mo R, Hui C-C, Henkelman RM, Nemer M, Black BL, Nagy A, Bruneau BG (2005) Tbx20 dose-dependently regulates transcription factor networks required for mouse heart and motoneuron development. Development 132:2463–2474.

Tariq MA, Kim HJ, Jejelowo O, Pourmand N (2011) Whole-transcriptome RNAseq analysis from minute amount of total RNA. Nucleic Acids Res 39:e120–e120.

Thorvaldsdóttir H, Robinson JT (2013) Integrative Genomics Viewer (IGV): high-performance genomics data visualization and exploration. Briefings in ….

Toki S, Kawasaki H, Tashiro N, Housman DE, Graybiel AM (2001) Guanine nucleotide exchange factors CalDAG-GEFI and CalDAG-GEFII are colocalized in striatal projection neurons. J Comp Neurol 437:398–407.

Topark-Ngarm A, Golonzhka O, Peterson VJ, Barrett B, Martinez B, Crofoot K, Filtz TM, Leid M (2006) CTIP2 associates with the NuRD complex on the promoter of p57KIP2, a newly identified CTIP2 target gene. J Biol Chem 281:32272–32283.

Topham MK, Prescott SM (2001) Diacylglycerol kinase zeta regulates Ras activation by a novel mechanism. J Cell Biol 152:1135–1143.

Trapnell C, Roberts A, Goff L, Pertea G, Kim D, Kelley DR, Pimentel H, Salzberg SL, Rinn JL, Pachter L (2012) Differential gene and transcript expression analysis of RNA-seq experiments with TopHat and Cufflinks. Nature Protocols 7:562–578.

Van Hook MJ, Wong KY, Berson DM (2012) Dopaminergic modulation of ganglioncell photoreceptors in rat. European Journal of Neuroscience 35:507–518.

Wong KY (2012) A retinal ganglion cell that can signal irradiance continuously for 10 hours. J Neurosci 32:11478–11485.

Xu J, Xiao N, Xia J (2010) Thrombospondin 1 accelerates synaptogenesis in hippocampal neurons through neuroligin 1. Nature Neuroscience 13:22–24.

Xue T, Do MTH, Riccio A, Jiang Z, Hsieh J, Wang HC, Merbs SL, Welsbie DS, Yoshioka T, Weissgerber P, Stolz S, Flockerzi V, Freichel M, Simon MI, Clapham DE, Yau KW (2011) Melanopsin signalling in mammalian iris and retina. Nature 479:67–73.

Young RW (1985) Cell differentiation in the retina of the mouse. The Anatomical Record 212:199–205.

Zhang M, Xia H, Li X, Wang X, Dong Y, Zhang T, Yu H (2010) C1 domain mediates CalDAGIII localization to the Golgi. Mol Biol Rep 37:3481–3485.

Zhou H, Yoshioka T, Nathans J (1996) Retina-derived POU-domain factor-1: a complex POU-domain gene implicated in the development of retinal ganglion and amacrine cells. J Neurosci 16:2261–2274.

